# In which cases does it make sense to classify an individual into one population and what are the alternatives?

**DOI:** 10.64898/2026.09.26.754621

**Authors:** Julika Schmitz, Jana Naue, Lukas Hoffmann, Zoe Lange, Ferdinand Suchanek, Sabine Lutz-Bonengel, Holger Dette, Peter Pfaffelhuber, Carola Sophia Heinzel

## Abstract

During biogeographical ancestry (BGA) analysis, an unknown individual for whom ancestry informative markers (AIMs) were analyzed is often assigned to one of a few reference populations. For some individuals, such an assignment is not meaningful: the two parents of the individual may come from different populations, or the individual may come from a population not included in the reference database. In this study, we examined classification methods in combination with 104 AIMs from the VISAGE Enhanced Tool to provide one of the following three possible outcomes: (i) in-reference (IR), with an assignment to one of the reference populations; (ii) admixed (AD), with parents from two in-reference populations; and (iii) out-of-reference (OR), where the individual’s ancestry is not sufficiently represented by the reference populations. We therefore compared seven classifiers in combination with class-specific acceptance thresholds for the OR decision. The best-performing method according to our selection criterion was a naive Bayes classifier over pairs of populations, which we call the recent naive Bayes (RNB) classifier. Populations held out of the reference panel were correctly rejected in 84.9% of cases. We further tested the method on 800 individuals from seven additional groups not included in the reference panel. Only 29.4% of individuals from four non-admixed populations were rejected, resembling joint ancestry with reference populations. In addition, we investigated the prediction accuracy in a newly collected dataset of 28 individuals with self-reported ancestry by analyzing AIMs using massively parallel sequencing.

## 1 Introduction

In recent years, forensic DNA analysis has evolved beyond the traditional identification of individuals using short tandem repeat (STR) profiles. In addition to individual identification, it increasingly includes methods for predicting externally visible characteristics, chronological age, and biogeographical ancestry of an unknown individual who left a trace at a crime scene or an unknown deceased person (Kayser and de Knijff, 2011; Kayser et al., 2023; Phillips, 2015; Schneider et al., 2019). Such extended analyses become particularly important when the determined individual STR profile does not match a (suspect) reference sample or any entries in (inter-)national DNA databases (McNevin, 2020; Samuel and Prainsack, 2019). In these cases, additional phenotypic or population genetic information can support investigations by narrowing down the pool of potential suspects or prioritizing investigative measures (Butler et al., 2011; Kayser et al., 2023; Samuel and Prainsack, 2019; Schneider et al., 2019). In Germany, for example, the prediction of eye, hair, and skin color as well as age is permitted under Section 81e of the Code of Criminal Procedure (regulating forensic DNA analysis), while the determination of biogeographical ancestry (BGA) continues to be the subject of social and legal debate. (Fleckhaus et al., 2025; Zieger, 2022). Methodologically, these prediction approaches rely on different genetic and epigenetic markers. Chronological age is typically predicted from age-correlated DNA methylation patterns at specific CpG sites (Freire-Aradas et al., 2017; Naue, 2023; Vidaki and Kayser, 2018), whereas externally visible characteristics and biogeographical ancestry are mainly predicted using single nucleotide polymorphisms (SNPs) (Kayser et al., 2023; Terrado-Ortunõ and May, 2024). For biogeographical ancestry prediction, ancestry informative markers (AIMs), typically biallelic SNPs showing substantial differences in allele frequencies between populations, are compared with reference datasets to infer an individual’s most likely population of origin (Phillips, 2015; Shriver et al., 1997; Kayser, 2015; Kosoy et al., 2009; Mogensen et al., 2020; Pereira et al., 2012, 2020; Rajeevan et al., 2012, 2020; Tvedebrink et al., 2017, 2018). BGA can be traced back to continental populations using forensic AIMs panels, including the populations of sub-Saharan Africa (AFR), Western Europe (EUR), Middle East (MEA), South Asia (SAS), East Asia (EAS), Native Americans (AMR), and Oceania (OCE) (de la Puente et al., 2021; Diepenbroek et al., 2020; Eduardoff et al., 2016; Kidd et al., 2014; Pereira et al., 2019; Resutik et al., 2023; Tillmar and Kling, 2025; Truelsen et al., 2021; Wang et al., 2007; Xavier et al., 2020). A finer differentiation of geographically close populations usually requires the analysis of a larger number of AIMs . In forensic practice, however, only small amounts of DNA are often available, so smaller marker panels with particularly high population divergence are preferred in order to ensure sufficient sensitivity (Phillips and de la Puente, 2021; Resutik et al., 2026). Various classification and clustering methods are used for the statistical evaluation of such markers, including the naive Bayes classifier SNIPPER (Phillips et al., 2007), the machine learning method TabPFN (Heinzel et al., 2025), and the admixture model (AM) as implemented in STRUCTURE (Pritchard et al., 2000). These methods use reference datasets with population-specific allele frequencies to calculate the probability of a DNA profile belonging to different groups of origin (Santos et al., 2016). The classification of individuals without mixed ancestry at the continental level is generally reliable (Escher et al., 2022; Kidd et al., 2014; Pfaffelhuber et al., 2020). However, more complex situations such as admixed ancestry or ancestry from populations that are not sufficiently represented in reference datasets cannot be handled correctly. Genetic admixture, in which different proportions of the genome originate from two or more populations, therefore poses a particular challenge. Hence, the question arises under which circumstances classifying an individual into a single population is justifiable. Two situations make it inadvisable. The individual may be admixed. In this setting, we have two possibilities: we can classify the parents of the individual into two different classes or, if this is not possible, classify the individual as “admixed”. Another reason why classifying an individual into a single population might not be meaningful is that the individual may originate from a population that is not represented in the reference data. A usable method must be capable of distinguishing both scenarios from those in which a classification is appropriate. Furthermore, this must be achievable without prior knowledge of the specific class category.

Admixed ancestry has been approached from several directions. STRUCTURE and ADMIXTURE answer it indirectly (Pritchard et al., 2000; Alexander et al., 2009); Pfaffelhuber et al. (2022) refine this to the possibility to two admixed parents. Posing admixture as a hypothesis test is more recent. Smyser et al. (2024) test whether *L* ancestral populations contribute to a trace against *K* := *L* + 1 in the alternative. Dette et al. (2026) test that the individual descends from exactly one reference population against the admixture alternative, using the constraint based bootstrap approach.

Tvedebrink et al. (2018) addressed the second situation. They test *H*_0_: the individual originates from population *k*, against *H*_1_: the individual is his or her own population, i.e. the reference dataset is not sufficient (referred to as “TVE” in this study). Tvedebrink and Eriksen (2019) extended the test to a pair of populations, so that a first-order admixed individual becomes the hypothesis (referred to as “TVP” in this study); that extension answers both questions at once, which is why TVP but not TVE can name two parental populations in the comparison below. Their approach and the approach developed in this study are related but differ because we aim to construct a classifier (assigning the individual to a discrete class) rather than a statistical test.

Outside forensics, a classifier that may refuse to decide is long established. Chow (1970) derived the optimal error-versus-reject trade-off for known distributions; classification with a reject option has since received a statistical learning treatment (Herbei and Wegkamp, 2006; Bartlett and Wegkamp, 2008; Cortes et al., 2016), and is called selective classification (Geifman and El-Yaniv, 2017). What we call out-of-reference is, in that literature, an open-set or out-of-distribution problem (Scheirer et al., 2013; Hendrycks and Gimpel, 2017; Geng et al., 2021), and a common strategy is to combine a scalar score of fit with a threshold calibrated on held-out data (Esposito et al., 2021; Wu and Ye, 2009; Masana et al., 2018).

Here, we combine a classifier with an *acceptance rule*: the classifier assigns a class to every individual, and the acceptance rule decides whether that class is reported or rejected. The acceptance rule has two parts: a score, which ranks how well an individual fits the reference panel, and a threshold, which turns that ranking into a decision. Because an individual assigned to a small population and one assigned to a large one are not comparable on the same scale, the threshold is fitted per predicted class (population), on data held out for that purpose.

As an alternative to classifying an individual into one population, we propose to classify its two parents into potentially different populations. This is related to the recent admixture model (RAM), where the aim is to estimate the share of the genome originating from each population for each parent (Pfaffelhuber et al., 2022); we therefore call the classifier the recent classifier. Because its classes are pairs of populations, a non-admixed individual is the pair (*k, k*) and admixture is a hypothesis of the model rather than a class that has to be learned from simulated training data. In total, seven classifiers for IR/AD/OR classification were considered in this study: naive Bayes (NB) (Phillips et al., 2007; Pfaffelhuber et al., 2020), recent naive Bayes (RNB), which is introduced here, the admixture model (AM) (Pritchard et al., 2000; Alexander et al., 2009), the recent admixture model (RAM) (Pfaffelhuber et al., 2022), TabPFN, a tabular foundation model (Hollmann et al., 2023, 2025), which Heinzel et al. (2025) applied to biogeographical ancestry, and the goodness-of-fit statistics classifiers (TVE, TVP) (Tvedebrink et al., 2018; Tvedebrink and Eriksen, 2019). Furthermore, an acceptance/rejection rule for classification is proposed.

This manuscript is organized as follows. Section 2.2 to section 2.4 describe the reference data, the evaluation design, the seven classifiers we compared (see above), the acceptance rule, and the quantities we report. Section 3.1 onwards compares the seven classifiers using reference data with one population absent in training, and examines the resulting error patterns. Appendix A describes the full pipeline, including the alternatives that were run and rejected, and appendix B gives the full probabilistic models. Figure 1 visualizes the proposed methods and fig. 4 gives an overview of the evaluation of the approach. Finally, the developed approach was applied to 800 independent samples from Ruiz-Ramírez et al. (2023), and novel buccal samples from 28 individuals with self-reported ancestry.

**Figure 1:**
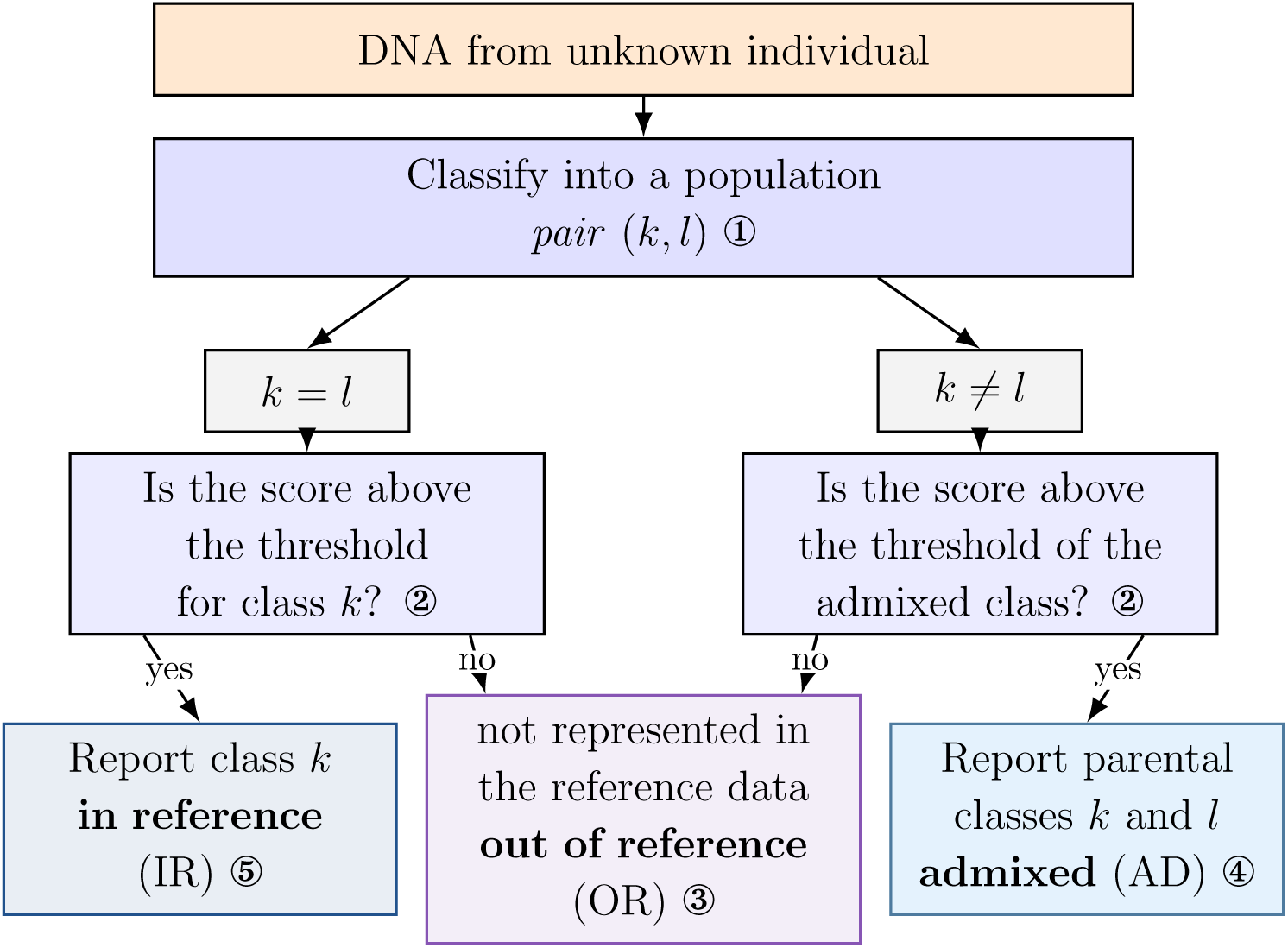
Overview of the classification pipeline and the quantities used for evaluation. For every pair-capable classifier (recent näıve Bayes (RNB), recent admixture model (RAM), admixture model (AM), and goodness- of-fit statistics classifier Tvedebrink and Eriksen (2019) (TVP)), we make a prediction of the population pair (*k, l*) ①. A score is then computed and compared with a threshold (which depends on the predicted class, with one shared threshold for Admixed) and the classification is retained only if the score is sufficiently high ②. If the score is below the threshold, the acceptance rule rejects the classification, resulting in out-of-reference ③. If the score is high enough and *k* ≠ *l*, the sample is reported as admixed ④. If the score is high enough and *k* = *l*, the sample is reported as in reference ⑤.

Every number and every figure below can be recomputed from the reference implementation at our GitHub repository. It contains the full code we ran on all samples. In addition, a standalone tool in that repository scores new samples under the recommended configuration and writes one report per sample.

## 2 Material and Methods

### 2.1 Workflow

An overview of the pipeline is provided in Figure 1. The inputs are the genotypes of the 104 AIMs of the VISAGE Enhanced tool (note that missing values are handled by the final model); see Section 2.2. For the BGA analysis, we apply a classifier (see section 2.3), and return one of three things for every sample:

**In-reference (IR)** The genetic data from the individual is most consistent with ancestry from one reference population.

**Admixed (AD)** The genetic data from the individual is most consistent with parents from two different reference populations;

**Out-of-reference (OR)** The population of the individual is not represented in the reference data.

No classifier directly returns “out of reference” as a class. In contrast, each classifier assigns an individual to one of its classes and provides a score for that assignment. If the score falls below a defined threshold, no final classification is reported because the assignment is rejected due to too high uncertainty. We call this the acceptance rule.

If we have *K* populations in the reference data, there is a total of 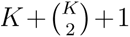 possible outcomes.

### 2.2 Used reference data

In this study, we used Supplementary Table S1A of Ruiz-Ramírez et al. (2023) as the reference dataset, which lists genotypes of the 104 AIMs of the *VISAGE Enhanced Tool for Appearance and Ancestry* for 4341 individuals in 14 groups (group names used as provided in Ruiz-Ramírez et al. (2023)). These markers also formed the basis for the analysis of 28 individuals in this study, allowing direct comparison (cf. section 2.7 and section 3.4).

We divided the 14 groups into two sets: Seven of them are continental populations and make up our reference panel, comprising *K* = 7 classes and *N* = 3541 individuals with the following composition in descending frequency: 935 (26%) East Asian (EAS), 827 (23%) European (EUR), 730 (21%) South Asian (SAS), 623 (18%) African (AFR), 240 (7%) Middle East (MEA), 109 (3%) Native American (AMR), and 77 (2%) Oceanian (OCE). Note that the sample distribution is highly imbalanced, both in class size and in genetic heterogeneity within classes, which can affect methods that weight individuals and populations differently.

The remaining seven groups, comprising a further 800 individuals, served as additional downstream test data, so they did not enter training and calibration of the classifiers; see section 3.3: central asian (64), east african (47), north african (29) and roma (3); middle east samples (’Sanger population data’) from (Almarri et al., 2021) (136); and two cohorts carrying an admixed label, admixed american (364) and admixed african (157). It has to be considered that two of the three possible BGA situations (AD and OR) cannot be observed directly in the reference data.

**Origin of the out-of-reference individuals** By definition, an out-of-reference individual is absent from the reference data. To construct this setting, we hold out one population at a time (we call this a *fold*) and treat its individuals as out-of-reference, with one modification: EUR and MEA are held out together. The two are at *F_ST_*= 0.047 (appendix A.1), the smallest separation in the panel, and holding out one of them alone most often leads to classification of these (out-of-reference) individuals into the other population. Appendix A.2 describes the alternative evaluation designs we ran as controls. This setting results in six different folds, depending on which population or populations are held out.

**Origin of the admixed individuals** There are no admixed individuals of known parentage in the reference panel, so that class is constructed in silico. In each run, admixed individuals are generated from the corresponding in-reference individuals. An offspring in the AD class is an F_1_ child of two in-reference individuals, one (independent) allele per marker from each parent, 15 per pair of parental populations, seeded identically for all methods. An out-of-reference individual is never used as a parent, so both parents of every admixed individual lie inside the reference. Note that the construction is restricted to first-generation admixture; no backcrosses or deeper admixture are introduced.

### 2.3 Classifiers compared for BGA prediction

Seven classifiers were compared. For each individual, each classifier returns a predicted class and at least one score; see (2.3) and appendix B.

**NB:** naive Bayes over populations (Phillips et al., 2007; Pfaffelhuber et al., 2020). Its classes are the seven populations plus one pooled AD class containing the simulated “training offspring”.

**RNB:** recent naive Bayes introduced here: naive Bayes over unordered population *pairs* with repetition, 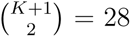 for *K* = 7, where a pair (*k, k*) is the non-admixed case. Because admixed individuals are represented directly within the model, no simulated admixed training offspring are required. Figure 2 illustrates the underlying model.

**Figure 2:**
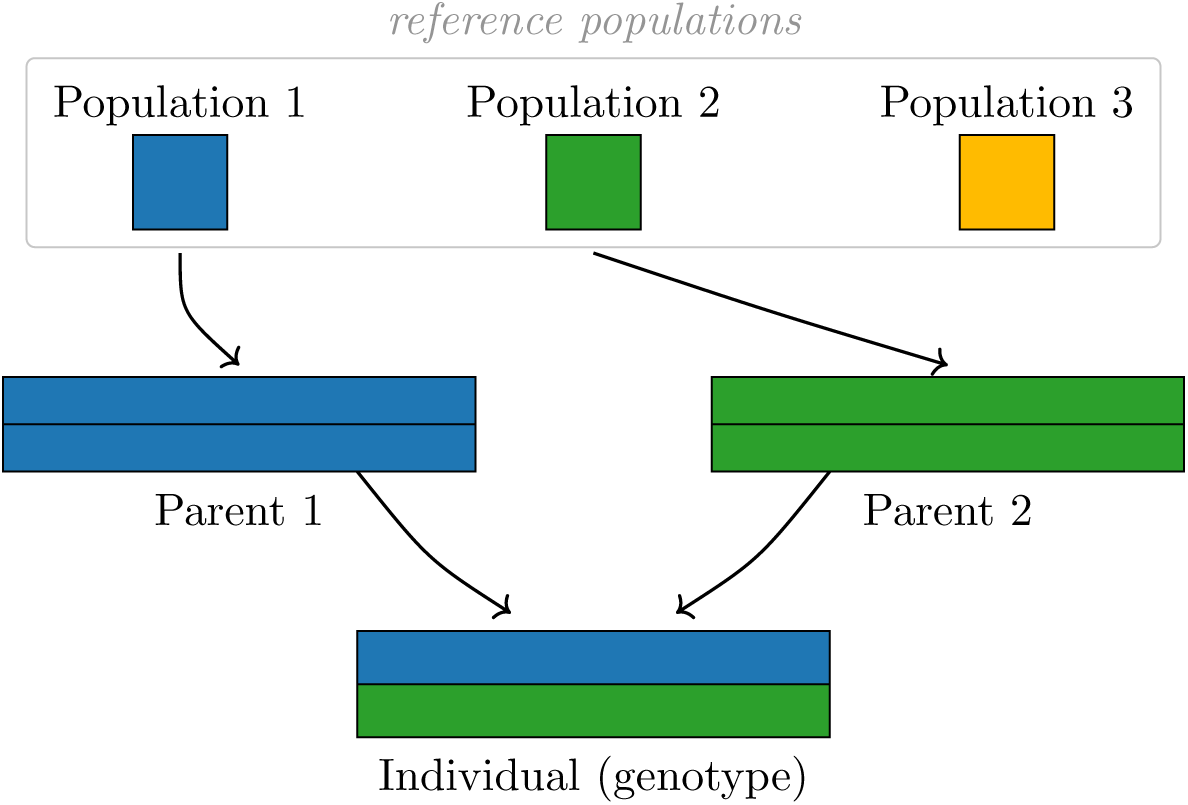
Model for the RNB. The three ancestral populations at the top are colour keys, shown as upright boxes; parental and individual genomes are represented by horizontal bars. Both parents are diploid and therefore carry two bars each. Under the RNB a parent belongs to exactly one population, so its two bars share one colour (in this example, population 3 contributes nothing). The individual receives one allele per marker from each parent, and the markers are inherited independently. The RNB scores the *pair* of populations directly, its estimate is therefore a pair (*k, l*), with the non-admixed case represented by (*k, k*) (both bars have the same colour).

**Figure 3:**
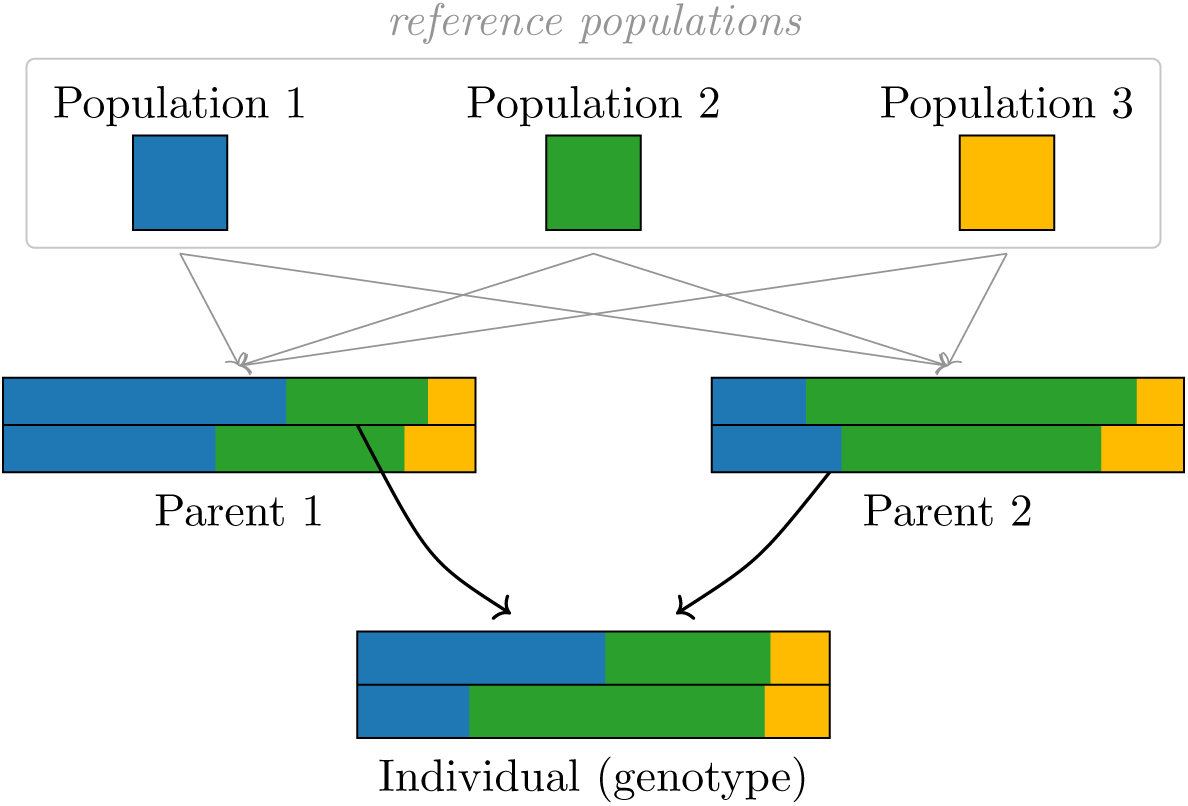
Model for the RAM. The same schematic as in fig. 2 but with already admixed parents: the parents are again diploid, but now each of their two bars is divided between the three ancestral populations rather than carrying one colour. Because the markers are inherited independently, the genomic contribution an individual receives from a parent is a mosaic drawn from both of that parent’s bars, and its composition is therefore that of the parent as a whole — which is what the individual’s two bars show here. Those two bars are the estimates *q*^(1)^ and *q*^(2)^, and this is why they are parental *halves*: each summarizes one parent, not one of that parent’s two bars. An individual is classified as non-admixed when the largest components of both halves agree and as admixed otherwise. See Pfaffelhuber et al. (2022) for more details.

**Figure 4:**
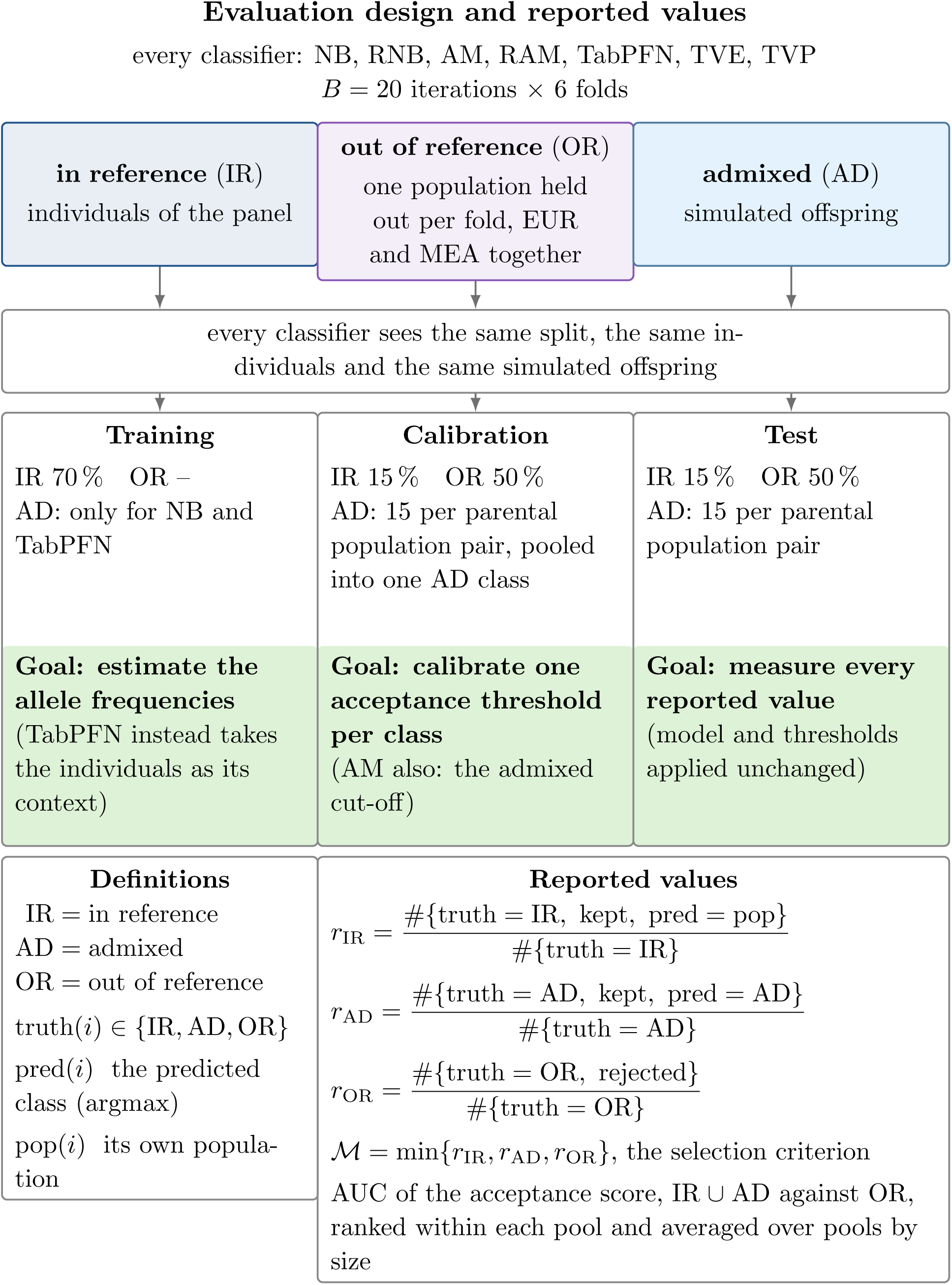
The evaluation design and the values reported from it. Every iteration has three groups of individuals — the panel itself (IR), the population held out in that fold (OR), and simulated offspring (AD) — and splits them into three parts, with separate subsets used to estimate allele frequencies, calibrate class-specific acceptance thresholds, and evaluate model performance. All classifiers are evaluated using the same split, individuals and simulated offspring.

**AM:** the admixture model (Pritchard et al., 2000; Alexander et al., 2009): continuous proportions *q* ∈ Δ*^K−^*^1^ (the *K* − 1-dimensional simplex) fitted per individual against the training data frequencies. It has no discrete non-admixed/admixed criterion, so we fit the criterion during calibration: an individual counts as admixed when its largest proportion falls below a cut-off.

**RAM:** the recent admixture model (Pfaffelhuber et al., 2022) (fig. 3): proportions fitted separately for the two parental halves. The individual is non-admixed when the two halves agree and admixed otherwise, so unlike AM the criterion needs no cut-off.

**TabPFN:** a pre-trained in-context tabular neural network (transformer) for tabular data (Hollmann et al., 2023, 2025), applied to biogeographical ancestry by Heinzel et al. (2025), fitted on the training data and giving a posterior over the classes it was trained on. Note that no likelihood-based probabilistic interpretation is available because the model does not provide a likelihood.

**TVE, TVP:** goodness-of-fit statistics, per population (TVE, Tvedebrink et al., 2018) and per population pair (TVP, Tvedebrink and Eriksen, 2019). The first has no notion of two parents and therefore never gives an admixed label.

**Restrictions of the classifiers** The pipeline must determine three aspects from the genotyped AIMs: the population assignment, whether the individual is admixed, and, if so, the contributing populations. Of the seven classifiers above, four can answer all three: RNB, RAM, AM and TVP. These are exactly the four that name a pair rather than a single class, and therefore the only ones for which an estimated pair is available. NB and TabPFN have a pooled admixed class but no notion of which populations contribute, so they answer two of the three. TVE has no admixed class at all.

### 2.4 Description of the procedure to evaluate the classifiers

#### Splitting of reference data

Each classifier described in section 2.3 was evaluated over 20 repeated iterations. Each iteration contains in-reference individuals as well as one or more out- of-reference populations (see Section 2.2). For the in-reference individuals, we have a 70*/*15*/*15 train/calibration/test split. Admixed individuals are simulated and out-of-reference individuals are defined as described above. In the calibration part, the offspring are pooled into a single AD class. All methods use identical splits, individuals and simulated offspring. The split is summarized in fig. 4. The composition of a split is the same in every iteration. Pooled over the six folds, one iteration’s test part contains on average 700 EAS in-reference individuals, 615 EUR, 546 SAS, 477 AFR, 177 MEA, 83 AMR and 60 OCE, together with 1275 admixed individuals. On the out-of-reference side a held-out population contributes 468, 414, 365, 312, 120, 55 and 39 individuals respectively. Within a single fold, MEA, AMR and OCE contribute fewer than 50 test individuals for evaluation each, which should be considered when interpreting population-specific results.

#### Training of classifiers

For six of the seven classifiers, allele frequencies are estimated from the in-reference individuals. TabPFN does not estimate allele frequencies at this step: it is an in-context learner, so the training individuals are handed to it as context and the pre-trained network uses them directly at prediction time. Two methods, NB and TabPFN (see section 2.3), additionally receive simulated admixed offspring here, because they can only predict a class they have seen; the other five obtain admixture structurally and train on the in-reference individuals alone.

#### Calibration

On the calibration set, every individual receives its acceptance score; see (2.3). Within each iteration, calibration predictions and scores from all six folds are pooled. For each classifier and acceptance score, one threshold *T_c,b_* is fitted within each predicted-class pool *c* in iteration *b*, by maximizing the mean of the class-specific retention and rejection rates defined in section 2.5. All admixed parental pairs share one Admixed pool and one acceptance threshold. The fitted thresholds are applied unchanged to the test predictions of the same iteration. For AM, a separate cut-off determining whether an individual is labeled Admixed is fitted on the calibration data within each fold, because the reference panel differs between folds.

#### Test of model performance

The thresholds estimated during calibration are applied unchanged to the test set. Every value reported below is computed on the test set.

#### Reported values

Two quantities (M and A) are reported for every comparison. Write truth(*i*) ∈ {IR, AD, OR} for the group an individual belongs to (in-reference, admixed, out-of-reference), pred(*i*) for the class with the highest score (also called the arg max) and pop(*i*) for an individual’s own population. Then

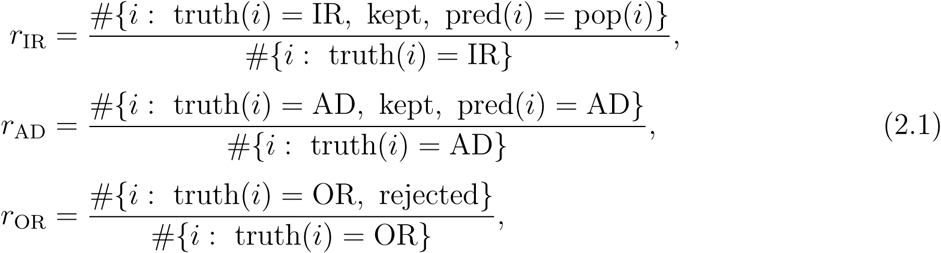

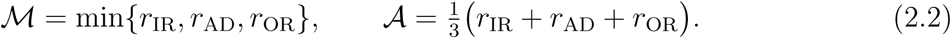

M is the selection criterion: a method is chosen by maximizing it over the classifier, acceptance score and threshold rule. The minimum requires the method to be adequate in every case. The accuracy A is the mean of the same three rates and is reported beside it. Note that *r*_IR_ requires the correct population label; assignment to a neighboring population is therefore counted as an error, as is rejection.

An additional model evaluation parameter is the area under the curve (AUC) of the acceptance score, computed with IR ∪ AD as positives and OR as negatives. The AUC measures the score’s ranking ability without requiring a threshold. We report it per pool, ranked inside each pool and averaged over pools by size. See Appendix A.6 for additional scores not reported here.

### 2.5 The acceptance rule: a score and a threshold

In standard (also called closed-set) classification, an individual is assigned to the class with the highest score. Here, motivated by open-set recognition and classification with a reject option, we extend this decision rule by introducing OR as an additional outcome (Scheirer et al., 2013; Hendrickx et al., 2024): the highest-scoring reference class is reported only if its score is at least the corresponding acceptance threshold; otherwise, the individual is assigned to OR.

Assume that *q_k_*(*x*) is the (a posteriori) probability of class *k* for individual *x*, and, if available, *P* (*x*|*θ*) the probability of obtaining individual *x*, which has *M* (*x*) successfully typed markers, under parameters *θ* (which depend on the classifier). This yields the two scores

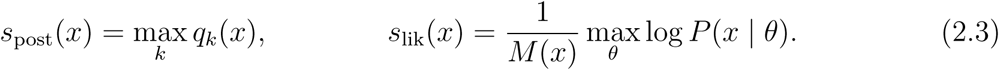

The available score depends on the classifier: TabPFN is discriminative and has only *s*_post_, NB, RNB, AM and RAM are full likelihood models and have both. TVE and TVP have neither: their statistic already measures fit in absolute terms. However, because a small value means a good fit here, we use its negative so that higher values indicate a better fit in all cases. Because 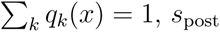, *s*_post_ cannot express that no panel class fits, whereas *s*_lik_ can.

Recall that a class is the set of individuals carrying that label as a truth. In contrast, we write pool *c* for the individuals a method assigned to class *c* by its argmax. Each pool receives its own threshold *T_c_*, fitted on the calibration set and applied unchanged to the test set.

A pool should keep individuals that belong to that label. Write correct(*i*) for the class individual *i* should be given: its own population if it is in reference and Admixed if it is admixed. An out-of-reference individual’s own population is not among the classes, so correct(*i*) ≠ *c* holds for it in every *c*: it has no correct in-panel label and therefore always counts on the rejection side. Because a high score indicates a good fit, define

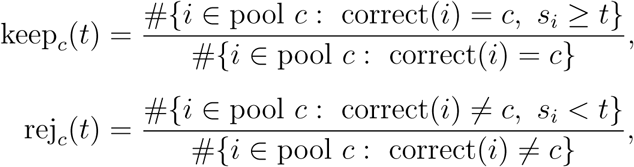

and the threshold *T_c_* maximizes <u>^1^</u> (keep*_c_* + rej*_c_*). Note that pool *c* is fitted only using data from pool *c*. Appendix A.4 gives two weaker criteria and additional alternatives not used in the reported analysis.

### 2.6 Using RNB on novel samples

While the training/calibration/test split had its purpose in recommending one method, we apply the RNB to novel samples below, and we have to adapt the pipeline for this. For deployment, RNB allele frequencies are estimated from all individuals in the seven reference populations. The fixed acceptance threshold for each predicted class *c* is obtained as the median of its iteration-specific calibration thresholds from the evaluation described in section 2.4, i.e. *T* ^deploy^ = median*_b_*_=1*,…,*20_ *T_c,b_.* Here, *T_c,b_* is fitted to the pooled calibration predictions from the six folds of iteration *b*, as described above. The resulting eight thresholds, one for each reference population and one for Admixed, are used unchanged for the additional 800 individuals and the 28 newly analysed samples (sections 3.3 and 3.4). A prediction is retained when *s*_lik_(*x*) ≥ *T* ^deploy^, where *ĉ*(*x*) is the predicted class.

### 2.7 Analysis of newly collected samples

#### Sample and library preparation

Buccal swab samples from 28 individuals were obtained with written informed consent using DNA-free swabs (Sarstedt, Nümbrecht, Germany). Approval was obtained from the Ethics Committee of the University of Freiburg (nr: 22-1427-S1). Biogeographical ancestry was inferred from information provided by the volunteers for the first (parents) and second (grandparents) generation; see table 5. DNA was extracted using the QIAamp DNA Mini Kit (Qiagen, Hilden, Germany) according to the manufacturer’s protocol and quantified with the Power Quant System (Promega, Madison, WI, USA) on a QuantStudio 5 System with QuantStudio Design and Analysis Software v1.5.2 (Thermo Fisher Scientific (TFS), Waltham, MA, USA). An implemented assay for analysis of BGA markers selected from the AmpliSeq Enhanced VISAGE panel was used for sample analysis. The modified assay used here consists of 106 autosomal and 83 Y-chromosomal markers with PCR primers designed for a multiplex assay (primer mix A and B) by Illumina for library preparation using the NEBNext technology (New England Biolabs (NEB), Ipswich, MA, USA) and was run on a MiSeq FGx (Verogen (now a Qiagen company), San Diego, CA, USA) . However, not all markers were used for BGA prediction within this study (cf. below). PCR reaction mix includes 1 ng of DNA, 5 µL Platinum Direct PCR Universal Master Mix (TFS), 5 µL Primer mix A or B, and DNA-free water to a volume of 20 µL. PCR was run under the following conditions: initial denaturation at 95 °C for 10 min, 35 cycles: 30 sec 95 °C (0.2 °C/s); 2 min 62 °C (0.2 °C/s); 2 min 68 °C, final elongation: 20 min 72 °C.

PCR products were purified with AMPure beads and library were prepared using the NEBNext UltraExpress DNA Library Prep Kit (E3325S, NEB) according to the manufacturer’s protocol. NEBNext Multiplex Oligos for Illumina (E6440S, NEB) were used for sample indexing. After the final clean-up, sample quality and quantity were determined on the TapeStation 4150 using the High Sensitivity D1000 DNA ScreenTape (both Agilent, Santa Clara, CA, USA).

#### Sequencing and data analysis

Samples were pooled equimolarly to prepare a 10 nM library. The final library, including 20 pmol phiX, was sequenced using 2×150 bp cycles (300 bp cartridge and micro or standard flow cell) on a MiSeq FGX (Verogen, now Qiagen). Paired-end fastq files were trimmed (quality score: 20) using Trim Galore v0.4.3/Cutadapt v2.0 and aligned to the GRCh38 (primary assembly) using BWA-MEM (release 0.7.17). SNPs were called and filtered using BCFtools 1.22 (mpileup, call, query) with a quality threshold of 20. For final SNP calling a coverage threshold of 30 was set and variants were visually verified using IGV.

#### Population assignment

To compare the classifier output with the participant information, the reported ancestry of the parents was mapped to the seven populations used in this study: AFR, AMR, EAS, EUR, MEA, OCE and SAS (consistent with the populations of the reference data). For example, Cambodian and Japanese reference individuals were assigned to EAS. Individuals of Philippine ancestry were mapped to EAS because Aeta, Agta, Luzon, Vizayan and most Igorot individuals in the reference table are assigned to EAS, although two Igorot entries are assigned to OCE in the reference data. In most cases, only country information was used for population inference; however, in some circumstances, additional available information (i.e. ethnogeographic information) was given priority over broad country labels. For example, Kurdish Iraq and Assyrian reference individuals are assigned to MEA, and unadmixed Peruvian and other Indigenous American populations are classified to AMR. For one individual (ID27), a tri-population target class (EUR/AMR/AFR) was assigned as its reported ancestry comprised three relevant components due to admixed parents.

Where the reference data contained heterogeneous assignments or no exact country match was available, the closest of the seven model populations was selected and the resulting ambiguity was retained for interpretation. For example, Iranian reference individuals occur under both MEA and EUR, whereas populations such as Ecuador, Chile, Costa Rica and Cuba are not represented directly. Ancestry reported as Turkey (Kurdistan region) is similarly ambiguous: generic Turkey and Turkish reference individuals are assigned to EUR, whereas Kurdish Iraq reference individuals are assigned to MEA. For samples specifically reporting ancestry from the Kurdistan region, the more specific Kurdish reference was prioritized and the respective ancestry was therefore mapped to MEA. Accordingly, these targets represent standardized approximations of the self-reported ancestry within the seven-class framework rather than independent genetic ground truth.

## 3 Results

### 3.1 Comparison of classifiers

#### Ranking of classifiers by M and AUC

Table 1 summarizes the seven classifiers under the evaluation design described in section 2.4: twenty iterations, six folds each, one threshold per population, fitted on the criterion of section 2.5. All values are computed on the test split only.

**Table 1:** Each method at its M-best acceptance score (see (2.2)), on the evaluation design of section 2.4: twenty iterations, six folds each, one threshold per population.

| Method | Score | $r_{\text{IR}}$ | $r_{\text{AD}}$ | $r_{\text{OR}}$ | $\mathcal{M}$ | $\mathcal{A}$ | AUC |
| --- | --- | --- | --- | --- | --- | --- | --- |
| **RNB** | $s_{\text{lik}}$ | 0.768 | 0.752 | 0.849 | **0.752** | **0.790** | 0.867 |
| RAM | $s_{\text{post}}$ | 0.744 | 0.766 | 0.740 | 0.740 | 0.750 | 0.788 |
| NB | $s_{\text{lik}}$ | 0.835 | 0.582 | 0.690 | 0.582 | 0.702 | 0.797 |
| AM | $s_{\text{lik}}$ | 0.783 | 0.832 | 0.559 | 0.559 | 0.724 | 0.744 |
| TabPFN | $s_{\text{post}}$ | 0.791 | 0.553 | 0.697 | 0.553 | 0.680 | 0.760 |
| TVP | either | 0.332 | 0.748 | 0.718 | 0.332 | 0.599 | 0.855 |
| TVE | either | 0.721 | — | 0.929 | 0 | 0.550 | **0.893** |

Based on the criterion M, the RNB classifier achieved the highest value with M = 0.752. RNB exceeded RAM by 0.012 in M and by 0.079 in AUC. The same ordering was observed across five seeds, with an average difference of 0.035; see table 4 in appendix A.8. TVE achieved the highest AUC. However, although TVE showed the strongest separation by AUC (0.893), it scores M = 0, because it never gives an admixed label.

#### Error rates

As a refinement of table 1, Figure 5 shows confusion matrices for all seven methods and the resulting error patterns. The remaining methods show distinct error patterns: TVP assigns 0.513 of the truly in-reference individuals to AD; NB classifies various admixed and out-of-reference individuals in the wrong classes; and AM classifies 0.370 of the out-of-reference individuals to AD, consistent with a limitation of an acceptance rule based on absolute fit. TVE has an empty AD column, because it cannot output the class, hence M equals zero. TabPFN rejects 0.412 of the admixed individuals outright. This is notable because an increased level of heterozygosity is characteristic for F1 admixed individuals (Garnier-Géré and Chikhi, 2013), and this signal is available to TabPFN during its in-context learning (defined as training here). RNB and RAM are the only two methods whose correct classification groups (IR/IR, AD/AD and OR/OR) all exceed 0.7. As mentioned before, RNB achieves a higher M than RAM, with all correctly labelled groups at ≥ 0.75 and a maximum of 0.21 for the true AD classified as OR incorrect group. RAM’s main weakness is the out-of-reference group, which are classified as in-reference in 0.243 of the individuals, in contrast to 0.080 for RNB.

**Figure 5:**
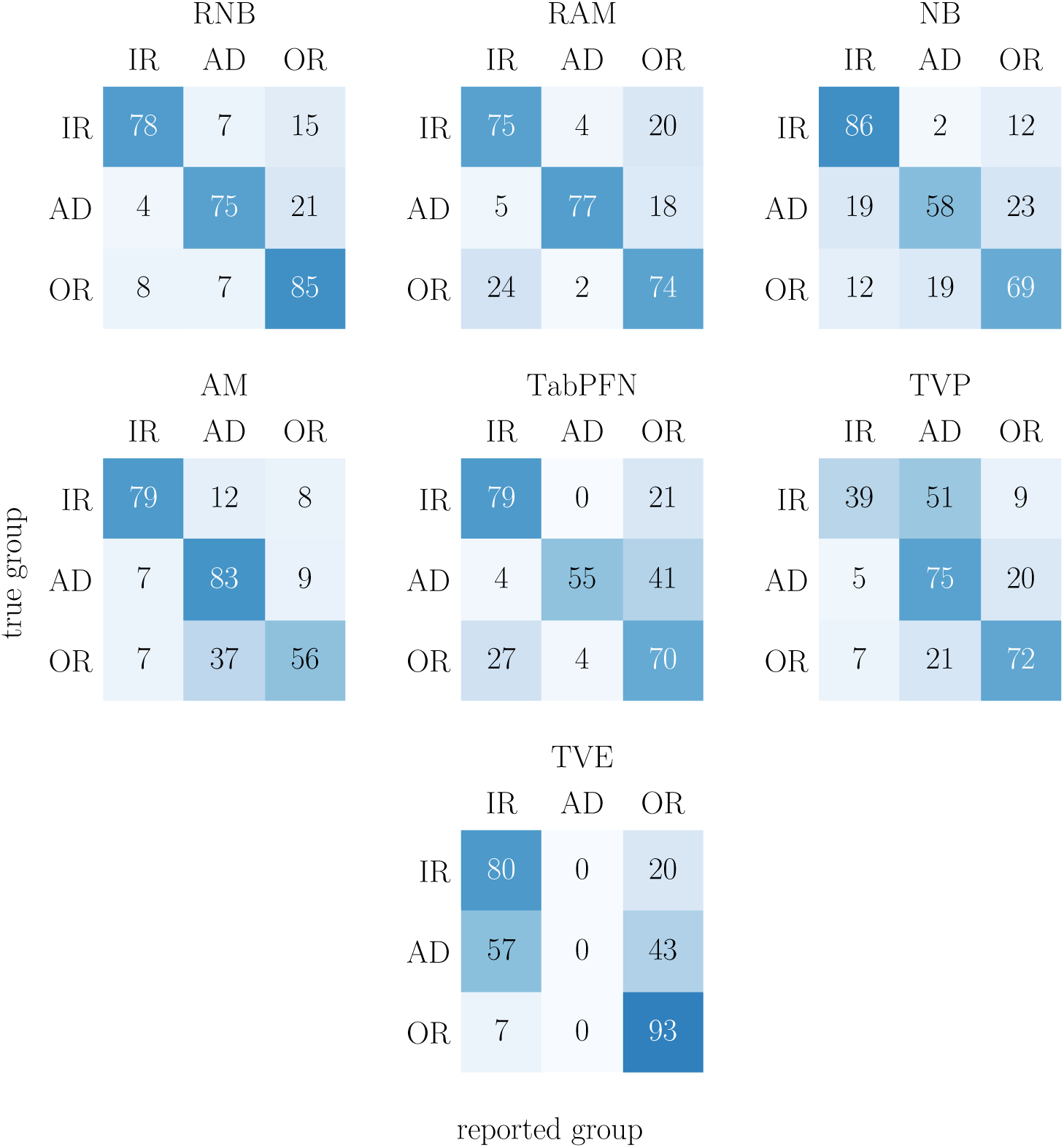
Confusion matrix showing the reported group (IR, AD, OR) in relation to the true group. Rows: true groups, columns: reported groups. Representation in percent of the row, so each row sums to 100 %. Here, the IR/IR cell counts individuals that were kept and not called AD, no matter which population was inferred (in contrast to *r*_IR_ in table 1), where the correct population must be inferred.

NB and TabPFN were excluded from subsequent analyses as their training on the admixed class led to poor performance, and both methods are unable to identify a specific pair of ancestral populations. TVE was also excluded from subsequent analyses because it cannot classify admixture.

### 3.2 The acceptance rule and the admixed class

Classification with a reject option is well established outside forensic genetics (Chow, 1970; Herbei and Wegkamp, 2006); within forensic ancestry inference it is rare, the exception being Tvedebrink et al. (2018); Tvedebrink and Eriksen (2019). Section 4 discusses our acceptance rule in relation to these approaches. Here, we examine the effect of the acceptance rule on RNB, the best classifier based on M (results for the other classifiers are in appendix A.9).

The left panel of fig. 6 provides a refined view of the RNB results shown in fig. 5: for in-reference populations, diagonal entries above the line show correctly classified individuals, whereas for out-of-reference populations, the final column below the line shows rejected individuals. AFR shows a high accuracy for both IR (93%) and OR (if AFR is left out, 100%) individuals, whereas some other populations show lower accuracy; e.g., in the MEA class 57% of in-reference individuals were kept, whereas 88% of the out-of-reference individuals were rejected. Note that in practice only the reported class is known, i.e. the column an individual is assigned to. As defined above, each column corresponds to a pool of individuals. Because columns are not normalized to sum to 100%, their composition can be examined separately. Here, the two weakest pools are MEA and OCE: of the individuals the acceptance rule keeps in the MEA pool only 58.1% are truly MEA, and in the OCE pool 62.5%; Section 4 reports the corresponding values for all eight pools.

**Figure 6:**
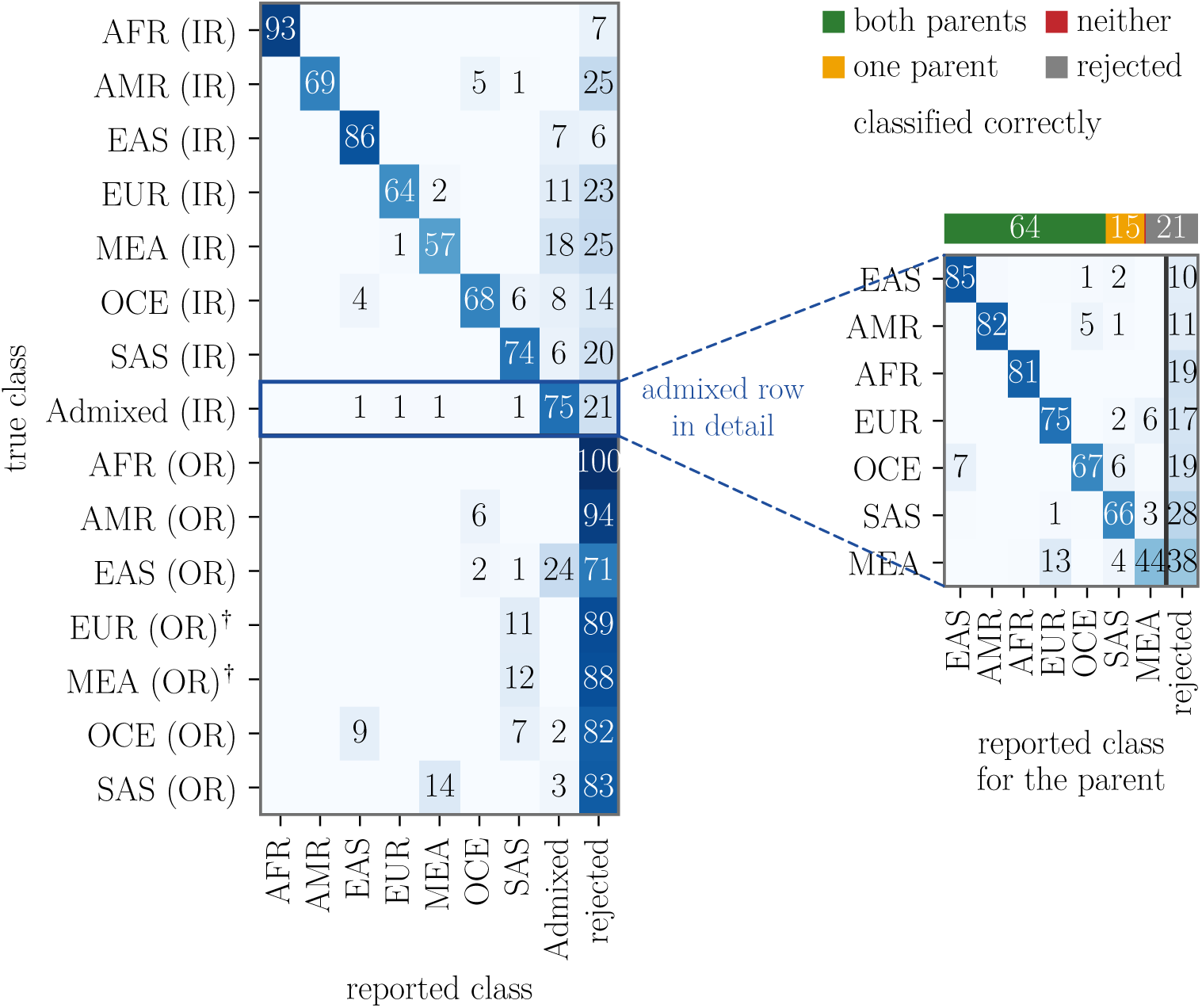
Performance of the RNB classifier after application of the acceptance rule. *Left:* Rows indicate the true class and columns the reported class; cells below 0.5 % are omitted and each row sums to 100 % (up to rounding errors). The upper rows show in-reference (IR) individuals from populations included in the panel. The lower rows show individuals from populations treated as out-of-reference (OR) in the corresponding fold; these samples should ideally be assigned to the rejected column. (Recall that EUR and MEA are held out together in a single fold (section 2.4) and are therefore marked with †.) *Right:* Detailed results for the true Admixed (IR) individuals. The horizontal bar summarizes the outcome per individual: both parental populations recovered correctly, one parental population recovered, neither recovered, or the individual rejected by the acceptance rule. The matrix below evaluates the same samples at the level of individual parental components. Each admixed individual therefore contributes two entries, one for each true parental population. Rejected individuals contribute both parental components to the rejected column. Both panels pool the six folds and twenty iterations, with the threshold in each fold and iteration fitted on the corresponding calibration set. For a given population, the IR row is based on the five folds in which that population is part of the reference panel, whereas the OR row is based on the fold in which it is held out.

The admixed individuals classified by RNB are examined in more detail (right part of fig. 6). This expands the Admixed (IR) row from the left panel: every individual of that row appears again, those the acceptance rule rejects in the rejected column and the rest at the parent pair they were given. Results are shown after applying the acceptance rule because 20.6% of these individuals are rejected and their parental assignments are therefore not reported by the pipeline. Of the two true parents, both are classified correctly in 64% of all cases and exactly one in 15%; both parents are incorrect in only 0.6% of cases. MEA shows the lowest parental recovery rate: 44% of its parents are recovered, 38% are rejected, and 13% are classified as EUR, which is the pair appendix A.1 measures at *F_ST_*= 0.047 and section 2.4 therefore holds out together. Individuals from SAS are also often rejected (28%) although the parents are well classified. The same analysis for the other three methods (RAM, AM, TVP) is in appendix A.9. These results depend on the use of M as the selection criterion. In addition, results for EUR and MEA depend on combining them for the out-of-reference class; see section 2.4. Additional evaluation designs were considered, and all seven methods were evaluated under each design. More details can be found in appendix A.3.

### 3.3 Individuals from groups the panel does not contain

The reference table of Ruiz-Ramírez et al. (2023) contains a further 800 individuals in seven groups that our reference panel does not contain (Section 2.2). Since none of them entered the training/calibration/test data, they can be evaluated directly using the fitted RNB configuration.

Four of the seven groups represent real populations absent from the panel, and only 29.4% of their individuals are rejected; compare with the rejection of 84.9% for the populations held out of the panel in the reported design (table 1). This difference is consistent with genetic distance to the reference panel: three of those four groups (north african, central asian, roma) lie close to a panel population (table 2), whereas the fourth (east african) has both the largest *F_ST_* and the highest rejection rate.

**Table 2:**
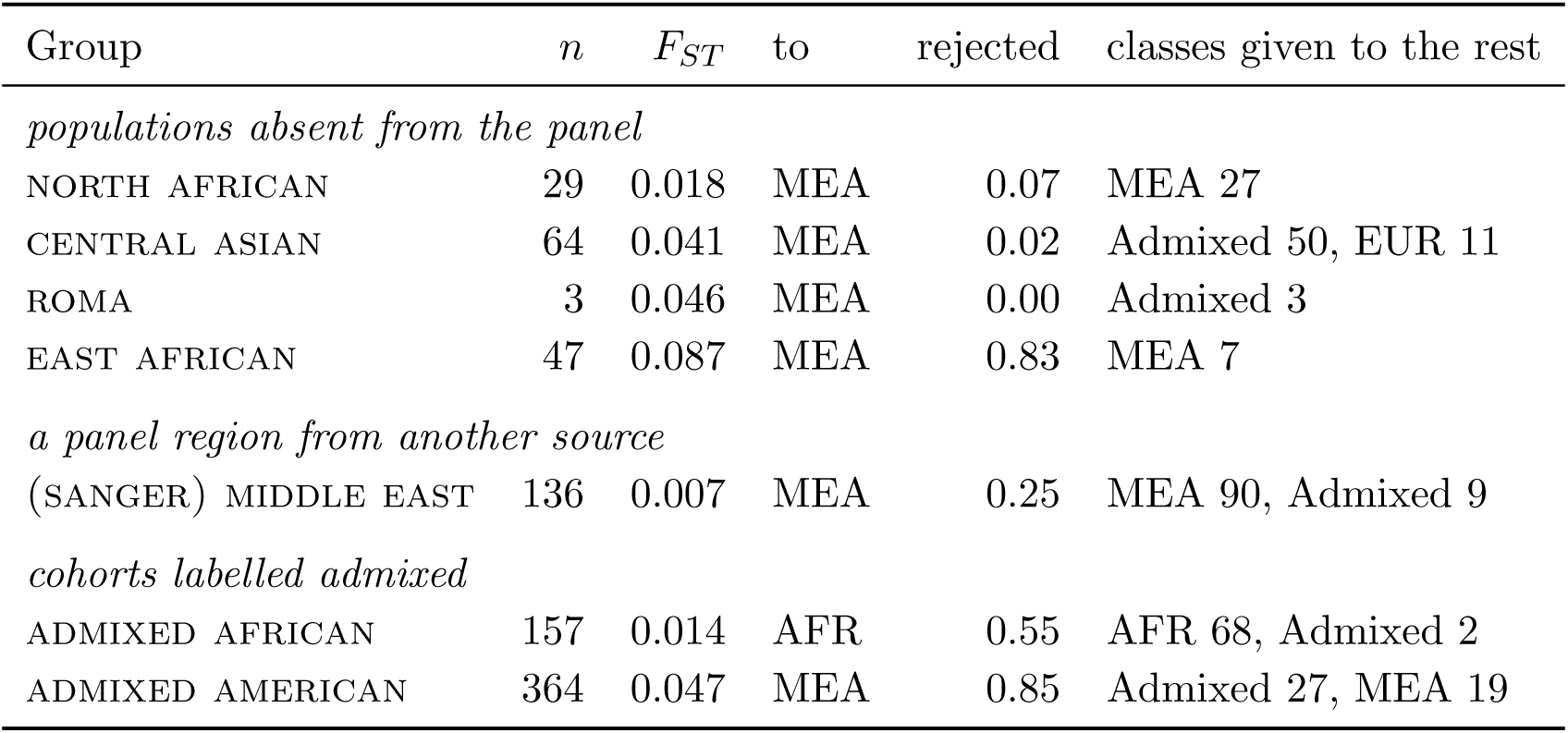
The additional 800 individuals of the reference table, scored by the deployed configuration using the RNB classifier. *F_ST_* of the corresponding group is computed against each of the seven panel populations with the estimator of appendix A.1, and the table reports the closest of the seven reference populations. The last column gives the classes that the individuals that passed the acceptance rule received most often. Groups are ordered by *F_ST_* within each block.

However, if an individual is not rejected, the reported class often provides a reasonable description relative to the reference panel: 27 of the 29 north african individuals are given MEA, and 50 of the 64 central asian ones are called admixed, most often as a mixture of EUR with MEA or with SAS. Appendix A.11 gives the remaining groups, among them the two cohorts carrying an admixed label, for which a correct answer is not defined.

### 3.4 Application to newly analyzed samples

#### Analysis of AIMs

All 28 individuals were analyzed using the modified panel comprising 106 autosomal markers. The marker rs12594144 (part of the 104 VISAGE Enhanced tool) was removed from further analysis because of insufficient coverage in most samples. Furthermore, rs9306481 and rs924381 were not considered for BGA prediction because they were not part of the SNP set used for model development. Thus, a maximum of 103 autosomal markers was available for BGA prediction. Amplification of rs12405776 slightly co-amplified an additional region (chr10:38330354–38330121). However, alignment against the complete reference genome allowed the amplicons to be assigned either to the rs12405776 locus or to the co-amplified region, enabling reliable variant calling of rs12405776. Due to the set quality threshold for final SNP calling, marker numbers ranged 78 to 103 (cf. table 5).

#### Classification using the RNB classifier and class-specific acceptance thresholds

All 28 individuals were classified with the recommended RNB classifier using the likelihood score *s*_lik_ and the class-specific acceptance thresholds described in section 2.5. That means a prediction was reported only when its likelihood score passed the threshold calibrated for the corresponding class. Samples that did not pass the relevant threshold were reported as *not classified*.

For evaluation, the reported population assignment was compared with the target derived from the participant information. (fig. 7, details in table 5). A single-population output *k* was treated as the pair (*k, k*). An *exact* match was defined if both populations were predicted correctly, whereas a *partial* match was assigned when at least one target component was classified correctly. No wrong classifications in respect to both parents were observed. Rejected samples were considered separately as *not classified* remaining an inconclusive result in a forensic case. ID27 was represented by three target populations but was rejected by the acceptance rule and therefore did not require an exact or partial pairwise classification.

**Figure 7:**
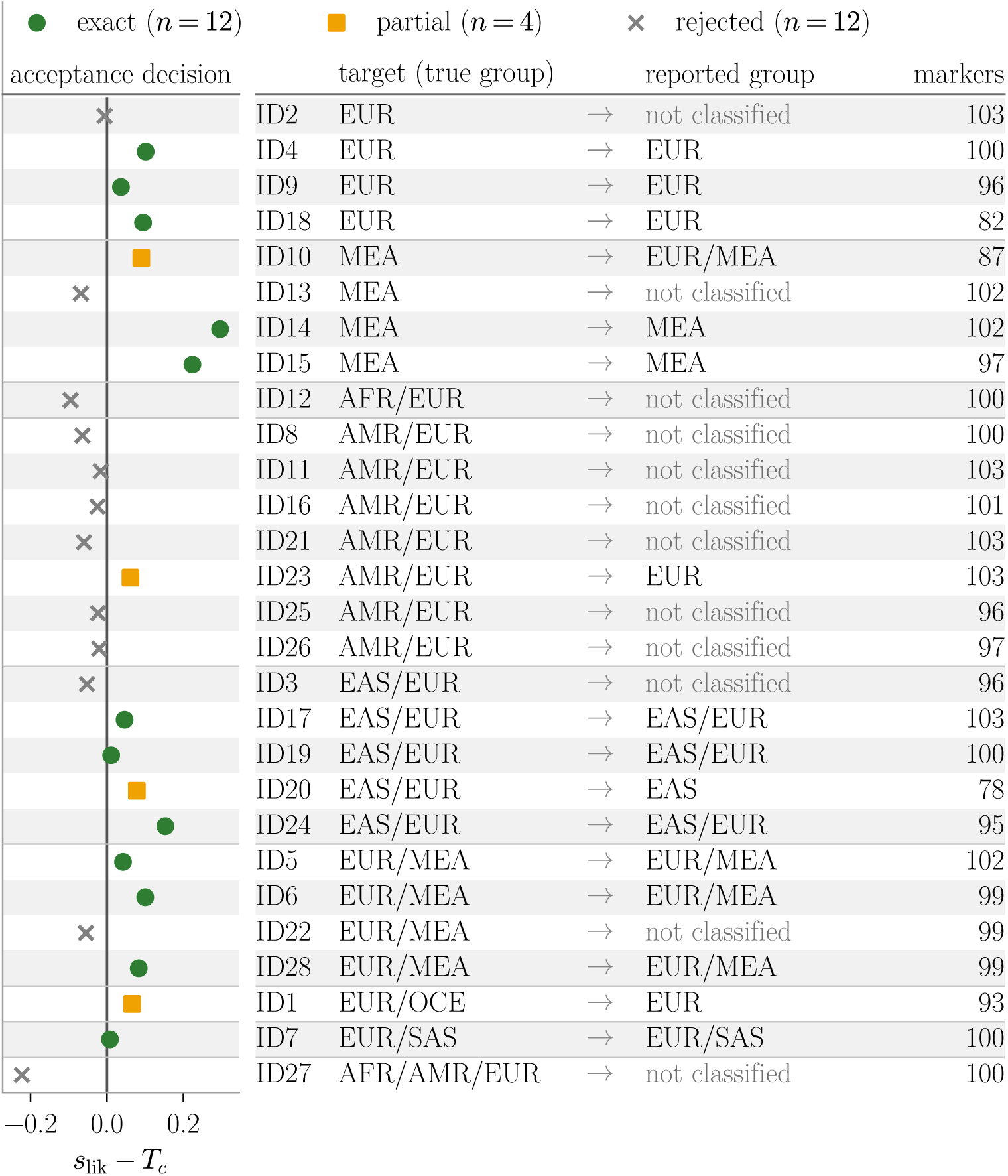
Classification results for the 28 empirical samples using the RNB classifier and the likelihood-based acceptance rule. The likelihood score relative to the class-specific acceptance threshold, *s*_lik_ − *T_c_*, is shown for each individual. Values at or above zero pass the acceptance rule, whereas values below zero result in *not classified*. Symbols distinguish exact, partial, and rejected classifications according to the ancestry targets derived from the participant information. For each individual, the target population assignment, reported classifier output, and number of markers available for classification out of the 103 usable markers are shown; the same data are listed in table 5. Rows are grouped by target, with single-population targets shown first, followed by two-population targets and the three-population target of ID27. Target populations were derived from the self-reported parental ancestry according to the population-group assignments used in the reference data. For two-population targets, the pair is unordered; a pair (*k, k*) is written as *k*.

Applying the RNB to the 28 individuals, 16 passed the acceptance rule and received a population assignment, whereas 12 were reported as *not classified* (fig. 7). Among the 16 reported classifications, 12 exactly matched the target and four were partial matches (one parent correctly assigned). Rejection and therefore a final non-classification occurred especially in cases of individuals of the AMR class (7 out of 8). This can be partly attributed to the fact, that at the parental level, also partly admixed ancestry is present. Admixed parents can lead to a non-evenly distribution of two populations within the child, e.g. in one of the rejected cases (ID 21), one parent is from Germany (EUR), whereas the other parent itself is an admixed parent from Italy (EUR) and Chile (AMR). Without using the acceptance threshold, only European was classified by the model, missing the American part of the second parent. In the case of ID27 a 3-population target AFR/AMR/EUR was present (parents: Portugal/Brazil (Indigenous) and Germany/Africa). As this is not represented by the model, a rejection occurred as expected (*s*_lik_ = −1.122, margin = −0.2227 ). (cf. table 5). The rejection of a classification is intended to avoid wrong classifications. Of all 12 rejected individuals, 9 individuals would have received a partial wrong classification (EUR was correctly classified in all cases), whereas 3 cases below the acceptance threshold would have had a correct assignment (ID2 EUR, ID3 EAS/EUR, ID16 AMR/EUR). In the case of the four partial classifications, in two cases the second component was missed (ID1, ID20) and in the other two cases, the individual was wrongly classified as being admixed without having a component (based on the self-reported ancestry) (ID10, ID23), respectively. ID1, with reported Australian and Scottish ancestry, was assigned EUR rather than EUR/OCE. Notably, the Australian reference data are themselves heterogeneous, with two Australian-labeled individuals assigned to OCE and one assigned to EUR. ID20, with German and Philippine ancestry (EUR/EAS) was classified as EAS only, whereas ID17 and ID19, with comparable German and Philippine or Cambodian ancestry, were both classified as EAS/EUR. ID10, for which both parental sides were reported as Turkey (Kurdistan region) (MEA only), was classified as EUR/MEA. Given that generic Turkish reference individuals are assigned to EUR whereas Kurdish Iraq is assigned to MEA, this result cannot be interpreted as a straightforward disagreement. In addition, EUR and MEA are the two closest reference populations in the panel, with *F_ST_* = 0.047. Finally, ID23, with German ancestry on one parental side and an Indigenous-American grandparent on the other, was mapped to AMR/EUR but classified as EUR only. Here, the AMR component was reported at the grandparent rather than parental level, whereas the RNB model represents each parent by one population.

As mentioned, the markers used for classification of the individuals varied. However, no simple relationship was observed between the number of missing SNPs and reduced accuracy (a partially correct result or rejection). Samples with comparatively large numbers of missing markers could still pass the acceptance rule: ID18 was classified exactly as EUR with 21 of the 103 markers missing, and ID20 was classified as EAS with 25 missing. Conversely, several samples with no missing markers, including ID2, ID11 and ID21, were rejected. Only successfully typed markers contribute to *s*_lik_, which is normalized by the number *M* (*x*) of available markers. The results therefore do not support marker dropout alone as an explanation for the rejected classifications.

## 4 Discussion

### A classifier with an admixed and a reject option

Under the reported design, we recommend the recent naive Bayes classifier with the likelihood score *s*_lik_; the method supports both admixed classifications and rejection. RNB achieved the highest selection criterion, M = 0.752, defined as the minimum of the accuracies for the in-reference, admixed and out-of-reference groups, and an AUC of 0.867. Several additional properties support this choice. For RNB, *s*_lik_ performed best across all four designs and both evaluation quantities. Its errors are the most evenly spread of all methods, with no off-diagonal cell above 0.21 (fig. 5). It names admixture structurally, as a hypothesis over pairs of populations the two parents come from, so it needs no simulated training offspring. Additionally, *s*_lik_ measures fit in absolute terms rather than as a posterior normalised over the panel, it can express that none of the panel populations provides an adequate fit. Finally, its estimate is closed form: it evaluates the likelihood of every possible pair and takes the largest, so the prediction is deterministic for a fixed input. However, this recommendation depends on the evaluation design, as the recent admixture model (RAM) performs better in some alternative designs. RAM under *s*_post_ achieves a higher criterion value in two of the three designs we do not report, leave-one-out without the EUR/MEA pairing and leave-two-out (appendix A.2, discussed in appendix A.3); in the first of the two it leads on the AUC as well. The recommendation is therefore conditional on the choice of reported design (appendix A.2), and would have to be revisited if that choice changed.

**Limitations** The approach has several limitations:

1. The admixed class in our calibration/test data consists only of simulated F1 hybrids. No admixed individual in the method comparison is real; this excludes other forms of admixture, including backcrosses and second-generation admixture.
2. The out-of-reference class depends on the genetic distance. Not surprisingly, a nearby missing population is not detectable as out-of-reference. Populations held out of the panel are rejected in 0.849 of cases, but these were distant from the remaining reference populations. (For this reason, we only left out EUR and MEA in combination.) The case Tvedebrink et al. (2018) raises, an individual from a population that the reference does not contain, is therefore a matter of degree.
3. In calibration, the thresholds are fitted under different panel sizes from those used for test data. While thresholds are fitted on a panel of 5 (if EUR and MEA are left out together) or 6 populations during calibration, they are used on 7 populations in the deployed system. The direction of that bias is known: as the number of panel populations increases, the best fit can only improve, so a threshold fitted at six under-rejects at seven.
4. Reference data for AMR (109 samples) and OCE (77 samples) are limited. Such small sample sizes may not represent the full heterogeneity of these populations and should be interpreted accordingly.

An additional limitation concerns the interpretation of predictions in casework. When analyzing a casework sample, the true ancestry is not known, but the reported class is. So, we should be interested in the question: provided we have a sample which is reported to be e.g. MEA, what is the probability that it really comes from MEA? (This is usually called the positive predictive value, PPV.) For the recommended configuration the question based on the pools is given in table 3.

**Table 3:** The PPV is the fraction of individuals which are true class *c* among those reported to be class *c*. (This is a reading down a *column* of fig. 6.) Take MEA, pooled over the six folds, an iteration keeps about 174 individuals in the MEA pool: about 100 are truly MEA, about 50 are South Asian individuals from the one fold in which SAS is held out of the panel — MEA and SAS are the second closest pair in the panel at *F_ST_* = 0.079 (appendix A.1) — about 12 are admixed and about 10 are European, which gives the 0.581.

| Pool | PPV | Pool | PPV | Pool | PPV | Pool | PPV |
| --- | --- | --- | --- | --- | --- | --- | --- |
| AFR | 0.999 | AMR | 0.959 | SAS | 0.816 | OCE | 0.625 |
| EAS | 0.977 | EUR | 0.964 | Admixed | 0.752 | MEA | 0.581 |

**Table 4:** Variation of the reported quantities across runs. The full comparison was repeated for five seeds, from 42 to 122 in steps of 20, such that no two runs use the same split. For each method, the first line gives the mean across the five runs and the second line gives the corresponding range. Every quantity is defined in (2.1) and (2.2); TVE has no admixed class, so its *r*_AD_ is empty rather than zero and its criterion is 0 by definition.

| | $r_{\text{IR}}$ | $r_{\text{AD}}$ | $r_{\text{OR}}$ | $\mathcal{M}$ | AUC |
| --- | --- | --- | --- | --- | --- |
| RNB | 0.770 | 0.762 | 0.843 | **0.760** | 0.877 |
|  | (0.764–0.779) | (0.752–0.776) | (0.836–0.849) | (0.752–0.766) | (0.867–0.888) |
| RAM | 0.752 | 0.765 | 0.727 | 0.726 | 0.786 |
|  | (0.737–0.764) | (0.758–0.772) | (0.709–0.743) | (0.709–0.740) | (0.782–0.789) |
| NB | 0.839 | 0.562 | 0.697 | 0.562 | 0.796 |
|  | (0.835–0.842) | (0.483–0.608) | (0.678–0.723) | (0.483–0.608) | (0.790–0.799) |
| AM | 0.786 | 0.837 | 0.554 | 0.554 | 0.743 |
|  | (0.781–0.791) | (0.827–0.853) | (0.543–0.562) | (0.543–0.562) | (0.734–0.751) |
| TabPFN | 0.787 | 0.545 | 0.690 | 0.545 | 0.761 |
|  | (0.784–0.791) | (0.528–0.553) | (0.677–0.698) | (0.528–0.553) | (0.755–0.766) |
| TVP | 0.330 | 0.748 | 0.718 | 0.330 | 0.856 |
|  | (0.327–0.333) | (0.724–0.765) | (0.702–0.740) | (0.327–0.333) | (0.854–0.858) |
| TVE | 0.732 | — | 0.928 | 0.000 | 0.895 |
|  | (0.721–0.738) |  | (0.925–0.933) |  | (0.893–0.898) |

**Table 5:**
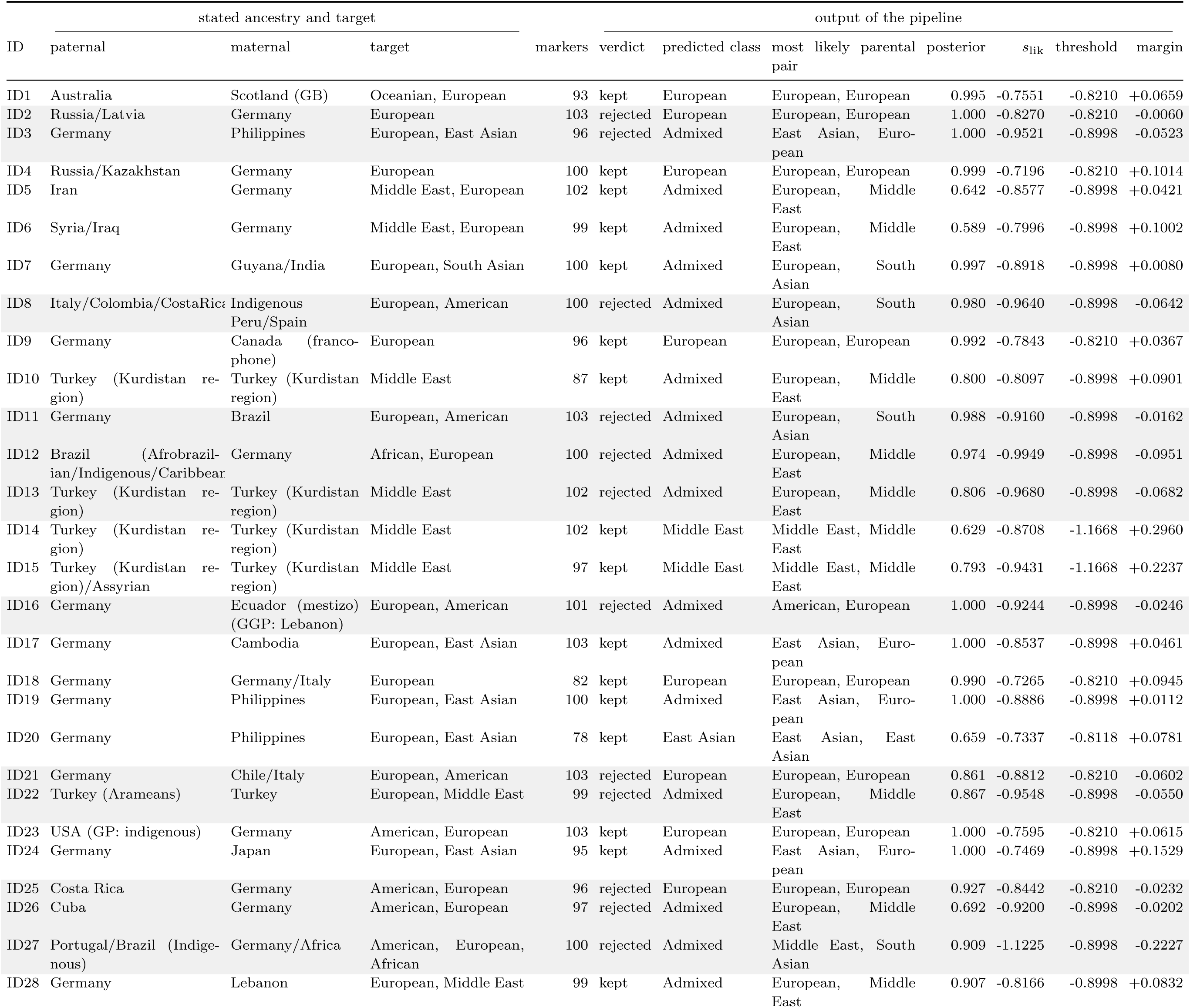
The 28 newly analysed samples, one row per individual. GP denotes grandparents and GGP great-grandparents. Grey rows were rejected by the acceptance rule; for these samples, the predicted class and parental pair indicate the most likely model output before rejection and are not reported as final classification results.

The same question can be put to the rejection: Given a sample is reported OR, how likely is it really OR (negative predictive value, NPV)? Pooled over the six folds and twenty iterations, an iteration rejects about 2168 individuals, of which about 1506 are genuinely out of reference, about 400 are in reference and about 262 are admixed. The negative predictive value is therefore 0.695, against an out-of-reference share of 0.311 in the test part.

Both, the positive and negative predictive value, depend on how common each group is, and in our study this is set by design and not by casework. In other words, for any real population, PPV and NPV will differ from the numbers just reported.

In our analysis, we used separate training, calibration and test sets within each split to avoid reusing the same observations for multiple stages. However, note that the selection criterion M is used twice: (i) to choose among several (39, to be exact) configurations formed by seven classifiers, two scores, and three threshold rules; (ii) as the performance we report for the chosen configuration. Consequently, the observed performance of RNB may overestimate what performance the selected configuration would achieve on fresh data (Varma and Simon, 2006; Cawley and Talbot, 2010, see also Kriegeskorte et al., 2009 for the general issue of non-independent selection and subsequent analysis). Yet another design decision was made after seeing the results: The joint EUR and MEA holdout was not fixed in advance. Rather, it followed from the comparison itself, where holding out either population alone led to its individuals being assigned to the other (section 2.4), irrespective of the classifier. This observation is corroborated by *F_ST_* = 0.047 for that pair against 0.079 for the next closest (appendix A.1). For these reasons, the recommendation of RNB should be interpreted as conditional on the reported evaluation design. We investigated the robustness of the resulting recommendation in two complementary ways. First, we repeated the complete comparison across five seeds with different random splits and RNB ranked first under M in every run. This demonstrates stability with respect to the random partition, but not with respect to the panel, since all five runs use the same 3541 individuals at the same 104 markers. Second, the selected RNB configuration was applied, without using either additional dataset for model selection or threshold calibration, to 800 held-out individuals (section 3.3) and 28 newly analyzed samples (section 3.4). These analyses provide additional out-of-sample evidence about the behavior of the deployed classifier, although neither of them yields an M-value. An evaluation separated from configuration selection could be obtained through nested resampling, with all configuration choices made within each outer development set, or through a further untouched evaluation dataset. Nested evaluation is constrained by the small number of reference populations and the limited sample sizes within some groups, with a single fold containing as few as 39 out-of-reference individuals. Additional independent evaluation would require suitable new data. In principle, following the simultaneous-inference approach to post-selection validity (Berk et al., 2013), confidence intervals covering the true M-values of all candidate configurations simultaneously would remain valid for the selected configuration. Although, applying this principle here would require a construction that accounts for our nonlinear criterion and dependencies in the evaluation design.

All analyses in this study are based on Supplementary Table S1A of Ruiz-Ramírez et al. (2023). Two properties of this dataset limit interpretation. The first is its composition. The seven labels are continental aggregates of very unequal size (from 935 EAS individuals down to 77 OCE). The labels are also heterogeneous within groups. Australian-labeled reference individuals are split between OCE and EUR, and individuals from Turkey are assigned EUR whereas individuals from the Kurdish region of Iraq are assigned MEA. These reference assignments affected the interpretation of the 28 samples, in ID1 and ID10 (section 3.4), where disagreement between the target and the reported group may reflect the reference assignment as well as classifier performance. The second property is the marker set. The 104 AIMs of the VISAGE Enhanced Tool were selected to differentiate continental groups, and the *F_ST_* matrix of appendix A.1 illustrates this limitation: EUR and MEA at *F_ST_* = 0.047 are not separable and have to be held out as one fold, and three of the four genuinely absent populations of section 3.3 lie closer to a panel population than EUR and MEA lie to each other, which helps explain why they are often not rejected. Consequently, the 104 AIMs can only answer *this population cannot be resolved by the panel* (rather than certain absence from the panel). A larger or differently chosen marker set moves that boundary, and how far it can be moved is an active question (Phillips and de la Puente, 2021; de la Puente et al., 2025; Resutik et al., 2026).

Resutik et al. (2026) is the study closest to ours: they put GENOGEOGRAPHER to 2820 individuals with a 1900-SNP panel over 21 metapopulations and – as we do – evaluate it on first-generation offspring simulated one allele per parent and marker. Enabling its admixture module, which tests all 210 pairs of reference groups, lowers the share of unassigned individuals from 3.79 % to 0.71 % and of discordant ones from 7.38 % to 1.06 %; but 63 % of the individuals are then assigned to between two and 21 of the 231 candidates at once and counted as concordant if the true one is among them. Remarkably, none of their 300 simulated offspring is left unassigned at all. We note that a classifier is the more practical instrument than multiple tests: a test returns every hypothesis it fails to reject, so its answer is a set that grows with the number of hypotheses tested and with any correction for multiple testing — which Resutik et al. (2026) note would be desirable here. A classifier returns one class by construction, so a report either gives one population, a pair of populations, or a rejection, which is the form casework needs.

#### The newly analyzed samples

Overall, the newly analyzed samples demonstrate both the applicability and the limitations of representing real-world ancestry using seven broad reference populations. When the acceptance rule was applied, the inferred population assignment was generally compatible with the standardized target derived from the participant information, and no accepted classification was completely discordant. At the same time, 12 of 28 samples were not classified, including individuals whose operational target populations were represented in the reference panel. However, the use of the acceptance rule avoided a partially incorrect classification in nine of the 12 samples.

ID27 illustrates the limitations of the two-population strategy particularly clearly. Its paternal ancestry was reported as Portugal/Brazil (Indigenous), whereas its maternal ancestry was reported as Germany/Africa, jointly comprising EUR, AMR and AFR components (cf. Results application). As mentioned, the sample was rejected by the acceptance rule. ID27 had the lowest likelihood score among the 28 empirical samples (*s*_lik_ = −1.122), consistent with the pronounced complexity of its reported ancestry and the difficulty of representing it within the model’s two-population framework. This case illustrates an inherent limitation of the RNB model for ancestry profiles that cannot be adequately represented by at most two parental populations.

A *not classified* result should therefore not be interpreted as direct evidence that an individual belongs to a population absent from the reference data. Rather, it indicates that the observed genotype did not provide sufficient support for reporting the class selected by the RNB classifier under the calibrated acceptance rule. Complex ancestry, geographically intermediate populations, ambiguities in the reference assignments and the necessary simplification of heterogeneous self-reported ancestry may all contribute to such outcomes.

#### The reject option

A classifier that may refuse to decide is long established outside forensics. Accordingly, the contribution of this paper is not the reject option itself but its adaptation to the BGA setting: a per-class rather than a global threshold, a likelihood score whose scale is comparable across pools of very different size, and a selection criterion, the minimum of the three rates, that prevents trading one outcome for another. Conformal prediction (Vovk et al., 2005) could provide a route to a stronger statement, of the form that the true population lies in a reported set with probability at least 1 − *α*. We did not pursue this approach here. The forensic approaches most closely related to our work are Tvedebrink et al. (2018); Tvedebrink and Eriksen (2019), implemented in GenoGeographer (Tvedebrink et al., 2017). These studies address the same setting considered here, an individual from a population the reference does not contain, and use a goodness-of-fit test of the profile against each claimed population, with the individual treated as its own population when nothing fits.

Our approach follows the same general principle: the quantity used for acceptance should measure fit in absolute terms rather than relative to the panel, motivating the use of *s*_lik_ as the score. Our approach differs in two respects. We construct a classifier rather than a statistical test, and we fit the cut-off separately for each predicted class on a calibration set rather than using a global test level. These differences are reflected in table 1, where TVE and TVP are included in the comparison: TVE is the strongest of all methods at rejecting out-of-reference individuals, at *r*_OR_ = 0.929, and among the weakest at naming the correct population, at *r*_IR_ = 0.721, while lacking an admixed class, while TVP names a population correctly for only 0.332 of the in-reference individuals. We find that the design using M as a selection criterion is more balanced, and identifies RNB as the recommended method.

## Data Availability

Due to data protection, underlying SNP data of the 28 individuals are not shown. They can be obained on reasonable request. The code to reproduce our results and to apply our methods to new data is available at https://github.com/pfaffelh/BGA-pairs-and-reject.

In particular, we provide a stand-alone Python script which produces a report on a novel genotype of an individual.

## Acknowledgments

The authors thank Timo Sänger for support with labwork.

## Funding

This work was funded by the Deutsche Forschungsgemeinschaft (DFG, German Research Foundation) – Project-ID 499552394 – SFB Small Data.

## Use of Generative AI

Large language models were used within the study and for manuscript preparatiopn. Anthropic’s Claude was used as a coding and writing assistant (wrote and ran parts of the evaluation and figure code, checked reported numbers against the pipeline output, and revised passages of the text). Furthermore, Google’s (DeepMind) Gemma 4 and OpenAIs GPT-5.6 Sol were used for improvement of phrasing . Every analysis, every number and every claim was specified, checked and approved by the authors, who take full responsibility for the content.

## Author Contributions

Conceptualization: Carola Heinzel, Sabine Lutz-Bonengel, Peter Pfaffelhuber Funding Acquisition: Sabine Lutz-Bonengel, Peter Pfaffelhuber, Holger Dette Investigation: Lukas Hoffmann, Jana Naue, Ferdinand Suchanek

Formal analysis: Carola Heinzel, Peter Pfaffelhuber, Julika Schmitz Software: Carola Heinzel, Peter Pfaffelhuber, Julika Schmitz

Validation: Carola Heinzel, Jana Naue, Peter Pfaffelhuber, Julika Schmitz

Writing -Original Draft: Carola Heinzel, Sabine Lutz-Bonengel, Jana Naue, Peter Pfaffelhuber, Julika Schmitz

Writing - Review & Editing: all authors

Supervision: Holger Dette, Carola Heinzel, Sabine Lutz-Bonengel, Jana Naue, Peter Pfaffelhuber

## Declaration of Conflicts of Interest

The authors declare that there are no conflicts of interest.

## Ethics approval

All procedures performed in studies involving human tissue were in concordance with the ethical standards of the institutional and/ or national research committee and with the 1964 Helsinki declaration and its later amendments or comparable ethical standards. Participants gave informed consent and approval was granted by the Ethics Committee of the University of Freiburg (22-1427-S1). This article does not contain any studies with animals performed by any of the authors.

## Clinical trial number

not applicable

## A More details on the pipeline

This appendix provides additional details on the design decisions underlying section 2.4 and section 2.5, including alternative approaches that were evaluated but not used for the reported analysis. The aim is to document the analysis sufficiently for the reported results to be reproduced.

### A.1 Divergence between the reference populations

Pairwise *F_ST_* values over the 104 markers were calculated using the Hudson estimator (Hudson et al., 1992), in its multiallelic form and as a ratio of averages over markers, as recommended by Bhatia et al. (2013), with the *n/*(*n* − 1) correction for within-population heterozygosity:

|  | AFR | AMR | EAS | EUR | MEA | OCE | SAS |
| --- | --- | --- | --- | --- | --- | --- | --- |
| AFR | — | 0.456 | 0.406 | 0.426 | 0.307 | 0.341 | 0.304 |
| AMR | 0.456 | — | 0.182 | 0.372 | 0.313 | 0.243 | 0.226 |
| EAS | 0.406 | 0.182 | — | 0.349 | 0.278 | 0.135 | 0.171 |
| EUR | 0.426 | 0.372 | 0.349 | — | **0.047** | 0.327 | 0.135 |
| MEA | 0.307 | 0.313 | 0.278 | **0.047** | — | 0.252 | 0.079 |
| OCE | 0.341 | 0.243 | 0.135 | 0.327 | 0.252 | — | 0.144 |
| SAS | 0.304 | 0.226 | 0.171 | 0.135 | 0.079 | 0.144 | — |

The pairwise *F_ST_* values range from 0.047 to 0.456, with a median of 0.278. These values are high in absolute terms, as expected because the 104 markers were selected for ancestry informativeness. Two pairwise comparisons are particularly relevant for the subsequent analysis. EUR and MEA show the smallest differentiation (*F_ST_*= 0.047) by a clear margin, followed by MEA and SAS (*F_ST_*= 0.079). The low differentiation between EUR and MEA motivates the paired hold-out fold described in section 2.4. AFR and AMR show the largest differentiation (*F_ST_*= 0.456).

### A.2 Three designs

We evaluated three designs for constructing the out-of-reference class:

#### D1, leave-*k*-out

Each fold removes *k* populations from the panel and treats their individuals as out-of-reference, so the reference panel in that fold contains 7 − *k* populations. An iteration runs all 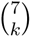 folds and pools their predictions. At *k* = 1 (*k* = 2), the same individual is treated as in-reference in six (fifteen) of the seven (twenty-one) folds and as out-of-reference in one (six), while its genotype remains unchanged.

#### D2, the reported design

D2 follows D1 at *k* = 1 with one modification: the fold that would hold out EUR and the fold that would hold out MEA are replaced by a single fold holding out both, resulting in six folds. The rationale is described in section 2.4. All other aspects of D1 remain unchanged.

#### D3, synthetic out-of-reference

The panel is not reduced: all seven populations remain available, and the out-of-reference individuals are generated synthetically. Both synthetic generators use the same basic step: for ancestor *k* and marker *m* with reference frequencies *p_km_*, a new population is drawn from the Balding–Nichols model,

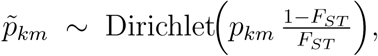

which reduces to the usual Beta distribution for two alleles and extends to multiallelic markers. Individuals are then generated under Hardy–Weinberg equilibrium from *p̃_km_*. We consider two variants: Mixed. For each individual, we select *m* ancestral populations, draw weights *q_i_* ∼ Dirichlet(*b, . . . , b*) for these populations, and form the mixed panel frequencies 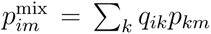. A Balding– Nichols step is then applied before generating one individual. Thus, out-of-reference individuals are generated from different simulated populations. With *m* = 7, mixtures lie at the centre of the simplex and are the easiest to detect, whereas with *m* = 2 the new population is derived from only two ancestral populations and is substantially more difficult to distinguish.

Drifted. One Balding–Nichols frequency vector is generated for each ancestral population, and all out-of-reference individuals assigned to that population are simulated from this vector. This variant isolates the effect of *F_ST_* on the divergence between the simulated population and its ancestor.

#### Reasons to report D2

The designs address different questions and should not be interpreted as interchangeable parameter settings. D2 evaluates performance on observed individuals from populations excluded from the reference panel, whereas D3 evaluates performance under a synthetic model whose results depend on the chosen simulation parameters. Across the mixture settings considered in D3, the accuracy ranges from 0.51 to 0.89, and the ranking of the methods can change. A result such as M = 0.75 for observed held-out populations therefore has a different interpretation from a value obtained under a synthetic generator. We therefore use D2 as the primary evaluation design. D3 is retained as a complementary analysis because it is the only design that keeps all seven populations in the reference panel, as in deployment, and the only design in which the degree of population novelty is a controlled variable.

### A.3 What the other designs show

The reported design holds out populations one at a time, with EUR and MEA held out together (section 2.4); appendix A.2 above gives the other constructions in full. All seven methods were evaluated under each of the four. This subsection summarizes why the remaining evaluation settings were not selected as the primary reported design.

#### The synthetic design

It leaves the panel complete and draws its out-of-reference individuals from an in silico model instead, with one allele-frequency vector per simulated population drawn from a Balding–Nichols distribution (Balding and Nichols, 1995); the construction is D3 above. We do not use this as the primary design because the degree and structure of novelty are imposed by the simulation rather than determined by observed populations.

#### Leave-two-out

Holding out two populations at once, rather than one, reduces the seven-population reference panel to five populations, removing nearly a third of the panel and therefore providing a substantially smaller reference than a single-population hold-out.

#### Leave-one-out without the pairing

As described above, EUR and MEA (*F_ST_* = 0.047, appendix A.1) are held out only together. Because these populations are closely related, holding out either one alone confounds the remaining population with the rejection option. Holding them out jointly reflects the effective resolution of the panel as six separable classes rather than seven, instead of evaluating a distinction that is not supported by the data.

#### The score

For the RNB, all four evaluated designs (D1 with *k* = 1 and *k* = 2, D2, and D3) favour the likelihood score *s*_lik_ over *s*_post_ for both reported quantities, with AUC differences of 0.12 to 0.28. In the reported design, it is 0.867 against 0.587 on the AUC and 0.752 against 0.533 on the M-criterion. For RAM, the second-ranked method, the two reported quantities favour different scores. Its AUC is higher for *s*_lik_ (0.834 versus 0.788), whereas its M-criterion value is higher for *s*_post_ (0.740 versus 0.482), a difference of 0.26. This disagreement between the two evaluation quantities provides no consistent basis for recommending one of the two scores for RAM.

### A.4 The threshold rule and the alternatives

#### Three criteria, nested

We considered three nested criteria for defining which individuals should be retained within a predicted pool: whether the predicted class is correct, whether the broader group is correct, or whether the individual belongs to the reference set at all. For an individual predicted as MEA: truth of an individual in the MEA pool

| Criterion | truth of an individual in the MEA pool |  |  |  |
| --- | --- | --- | --- | --- |
|  | MEA | other panel pop. | admixed | out-of-reference |
| **name** (in use) | keep | **reject** | **reject** | reject |
| coarse group | keep | keep | neither | reject |
| reference membership | keep | keep | keep | reject |

The distinction between these criteria is important. Under the coarse-group criterion, an inreference individual retained under the wrong population is still counted as correct, whereas an admixed individual assigned to a population pool contributes to neither side of the threshold objective. Consequently, retaining or rejecting such an individual does not affect the fitted threshold.

#### Three weightings

Within each pool, the two rates defined in section 2.5 are combined in one of three ways: weighting them according to the number of individuals on each side (name count, equivalent to the overall proportion correctly handled by the threshold), weighting them equally (name bal), or taking their minimum (name min). All three criteria combine the same two rates but assign different importance to imbalances between them. In particular, name count can achieve a high value at the expense of the smaller group, whereas name min is determined by the weaker of the two rates.

`Name_bal` is used throughout the reported analysis. Although `name_count` yields higher M-criterion values in some settings, by up to 0.15—more than most differences between methods— this occurs at the expense of individual predicted classes: it results in 94 of 2016 class–iteration combinations with no retained individuals, compared with none for `name_bal`, and results in no accepted predictions for the Admixed class for one method. Thus, `name_count` increases the overall criterion by leaving some classes without accepted predictions, which is precisely the behaviour the criterion was intended to avoid. `Name_min` is stricter still with respect to this imbalance and reduces performance on other quantities. Consequently, criterion values obtained with different threshold-weighting rules are not directly comparable.

### A.5 How sharply the threshold is determined

A fitted threshold is informative only if the objective it maximises has a sufficiently well-defined maximum. Figure 8 shows the two components of this objective as the threshold is varied for the reported configuration.

**Figure 8:**
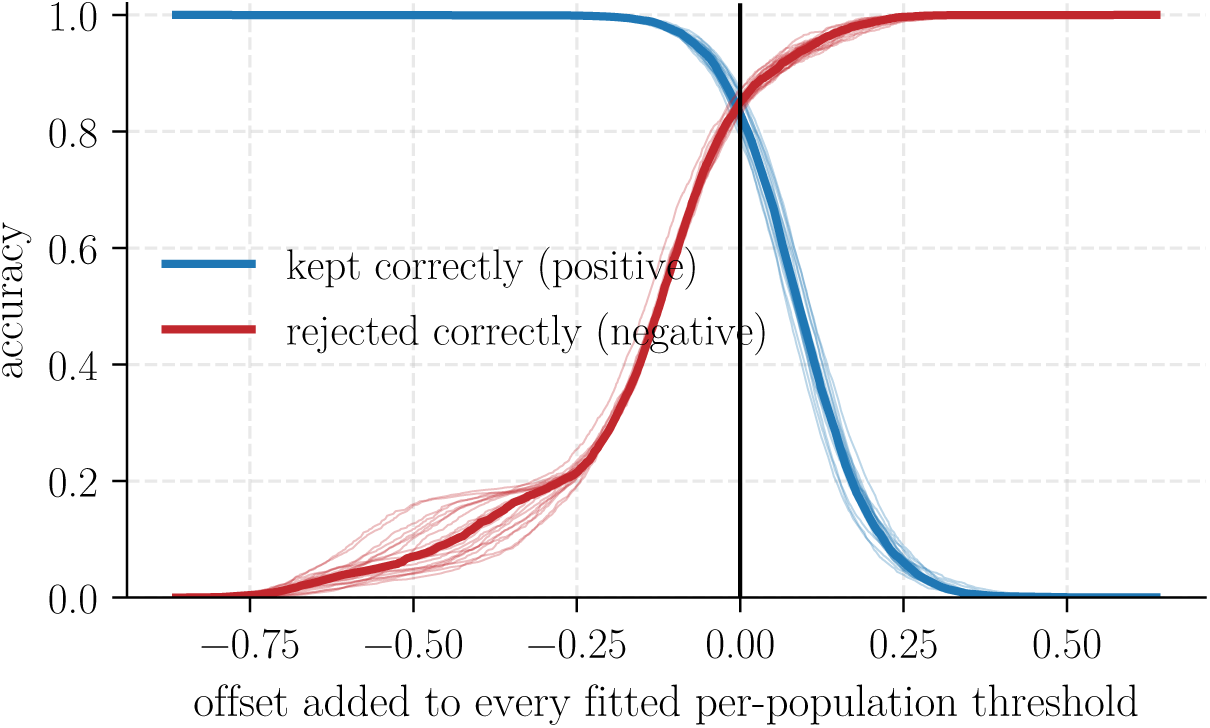
In each iteration, a class-specific threshold is determined. Each thin line corresponds to one of the twenty iterations, and the heavy line shows their median. The blue curve shows the proportion of individuals that should be retained and are retained, whereas the red curve shows the proportion that should be rejected and are rejected. On the *x* axis, the fitted threshold of each individual’s predicted class has been subtracted from its score. Zero (vertical line) therefore denotes the fitted operating point, and the axis represents a common offset from that point.

### A.6 Quantities that are not reported

Four additional quantities were computed but are not reported as primary metrics. Three are discussed here because they are natural alternatives to the quantities used in the main analysis.

Overall accuracy, with each individual weighted equally, depends on the class composition of the test set. A method that accepts nearly all individuals can therefore obtain a high value when the test set is dominated by individuals that should be retained. The accuracy A of section 2.4 is the same quantity with the three groups weighted equally.

Balanced accuracy for retained versus rejected individuals merges in-reference and admixed individuals into a single positive group and therefore does not evaluate whether the retained class label is correct. For example, an admixed individual retained under a population label and an in-reference individual retained under Admixed both count as true positives.

Accuracy over all nine outcomes – the seven populations, Admixed, and rejection – assigns equal weight to each outcome. Consequently, Admixed and rejection each contribute only one ninth of the total. Under the reported design, this weighting changes the ranking substantially and places a method with M = 0.124 in the middle of the comparison. More generally, a per-class mean across seven population classes, one admixed class, and rejection gives relatively little weight to the two outcomes that are central to the present pipeline.

Taking the minimum over populations is also problematic because, in practice, the lowest values largely reflect the trade-off between MEA and EUR, while three populations fall below the required *n* ≥ 50 sample-size threshold and would be excluded entirely. The resulting minimum would therefore be taken over only six of the nine possible outcomes. Population-specific behaviour is instead examined separately in fig. 6 and in the analysis below.

### A.7 Two checks the criterion cannot make

The global criterion does not capture either of the following checks for the same reason: small predicted pools contribute in proportion to their size. OCE, for example, contains 77 of the 3541 reference individuals, so poor performance for this population has only a small effect on the global criterion while potentially affecting the entire OCE pool.

A first additional check is whether any predicted class becomes unreachable after thresholding. The three global rates do not distinguish the weighting rules on this property. Across the eight output classes, however, the ordering is reversed: the weighting rules with the highest criterion values are those that leave classes with no accepted predictions. The separable, pool-specific weighting rules cannot fail in this way because each pool is fitted using its own observations.

A second check concerns the composition of the accepted part of each pool. Restricted to individuals retained by the threshold, prec*_c_* is the proportion whose correct class is *c*. This quantity describes the reliability of an emitted class label and is therefore directly relevant to casework interpretation. It is the precision counterpart to *r*_IR_, which instead describes what happens to an individual of known type. The values for the recommended configuration are tabulated in Section 4.

### A.8 How much the numbers move between runs

All values in section 3 are based on one run comprising twenty iterations with seed 42. The seed determines both the twenty train/calibration/test splits and, for the two admixture models, the initialization of the fitting procedure. To assess sensitivity to these random choices, the full comparison was repeated for five seeds separated by increments of twenty, such that no two runs use the same split.

RNB has the highest criterion value in all five runs, with margins of 0.012, 0.053, 0.048, 0.040, and 0.022 over the next method; its AUC margin ranges from 0.079 to 0.099. In the run reported in section 3, RNB has its lowest criterion value as well as its smallest criterion margin over RAM and its smallest AUC margin among the five runs. The reported run is therefore the least favourable of the five for the relative performance of RNB, so the main-text comparison understates rather than overstates its advantage.

Variation across seeds differs between methods. The two goodness-of-fit statistics and the two naive Bayes classifiers vary by at most two to three percentage points for most quantities, because their predictions are based on closed-form expressions and the seed primarily changes the allocation of individuals to the data splits. The main exception is the admixed rate of NB, which ranges from 0.483 to 0.608 because the admixed class is learned from simulated offspring and therefore depends more strongly on the particular draw. The recent admixture model shows its largest variation for the out-of-reference rate (0.709 to 0.743). As discussed in Section 4, its optimisation can converge to different optima depending on the initialisation.

### A.9 The acceptance rule and the parents, method by method

Figure 6 shows, for the recommended method, the outcome of the acceptance rule and, for predictions labelled Admixed, the inferred parental populations. The figures below show the corresponding results for the other six methods, using the acceptance score reported for each method in table 1 and following the order in section 2.3: first the three methods retained for detailed comparison alongside the recent classifier, followed by the three remaining methods. Figures 9 to 11 show both components; figs. 12 to 14 show only the left component because these three methods do not provide a parental-population pair. Rows, columns, the line separating in-reference from out-of-reference rows, and the dagger follow the conventions of fig. 6. The parental rows are shown in the same order, and every panel pools the same six folds and twenty iterations, allowing direct comparison across methods and with the main-text figure.

**Figure 9:**
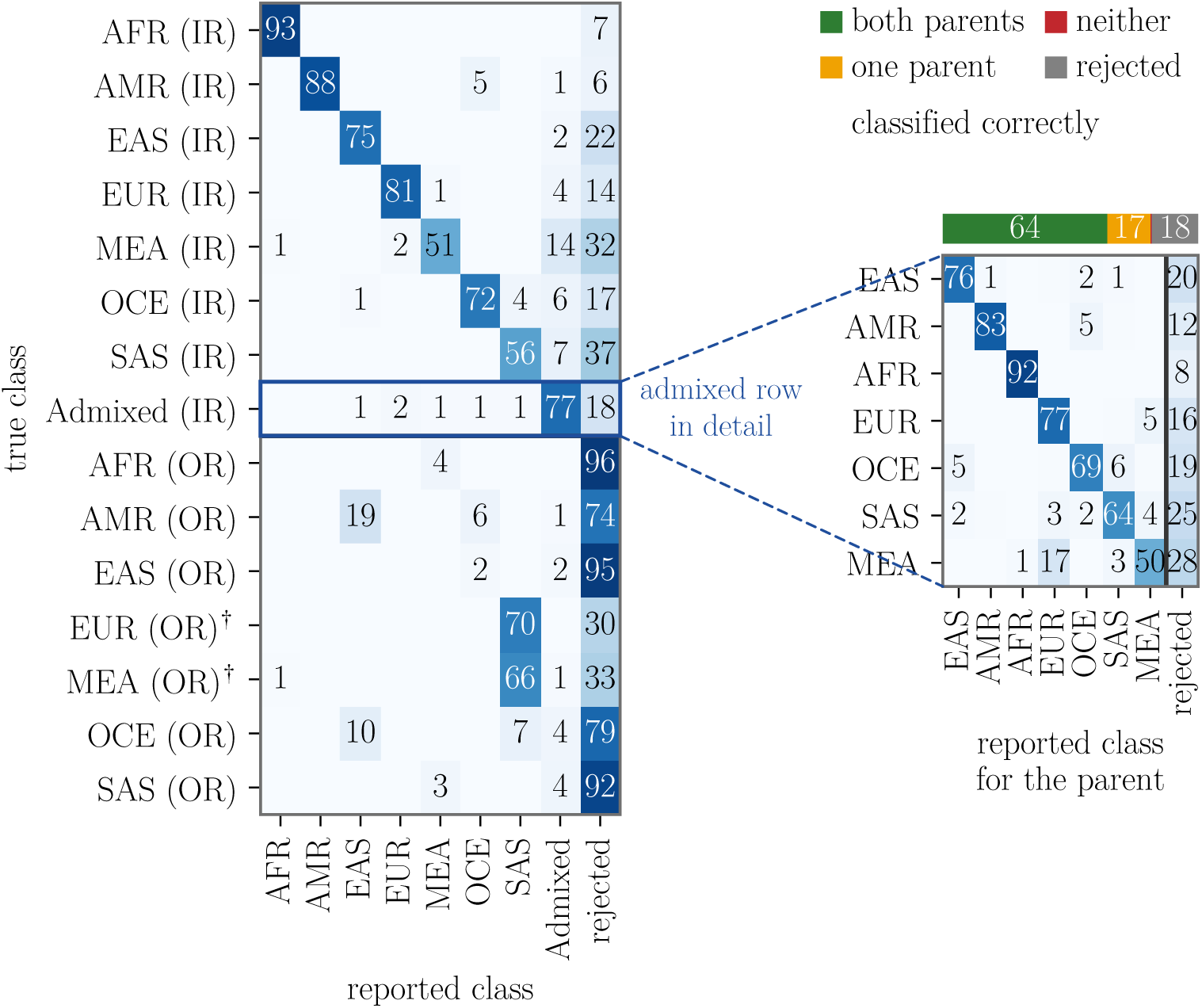
The recent admixture model under *s*_post_. Among the six comparison methods, its decision pattern is most similar to that of the recommended RNB classifier in fig. 6, consistent with RAM ranking second on the criterion. For out-of-reference individuals, RAM assigns a population label in 0.243 of cases, approximately three times the corresponding proportion for RNB, resulting in a lower rejection rate. For parental-population recovery, both parents are recovered in 0.645 of cases compared with 0.636 for RNB, while 0.182 are rejected compared with 0.206 for RNB. MEA is the weakest parental row for both methods.

**Figure 10:**
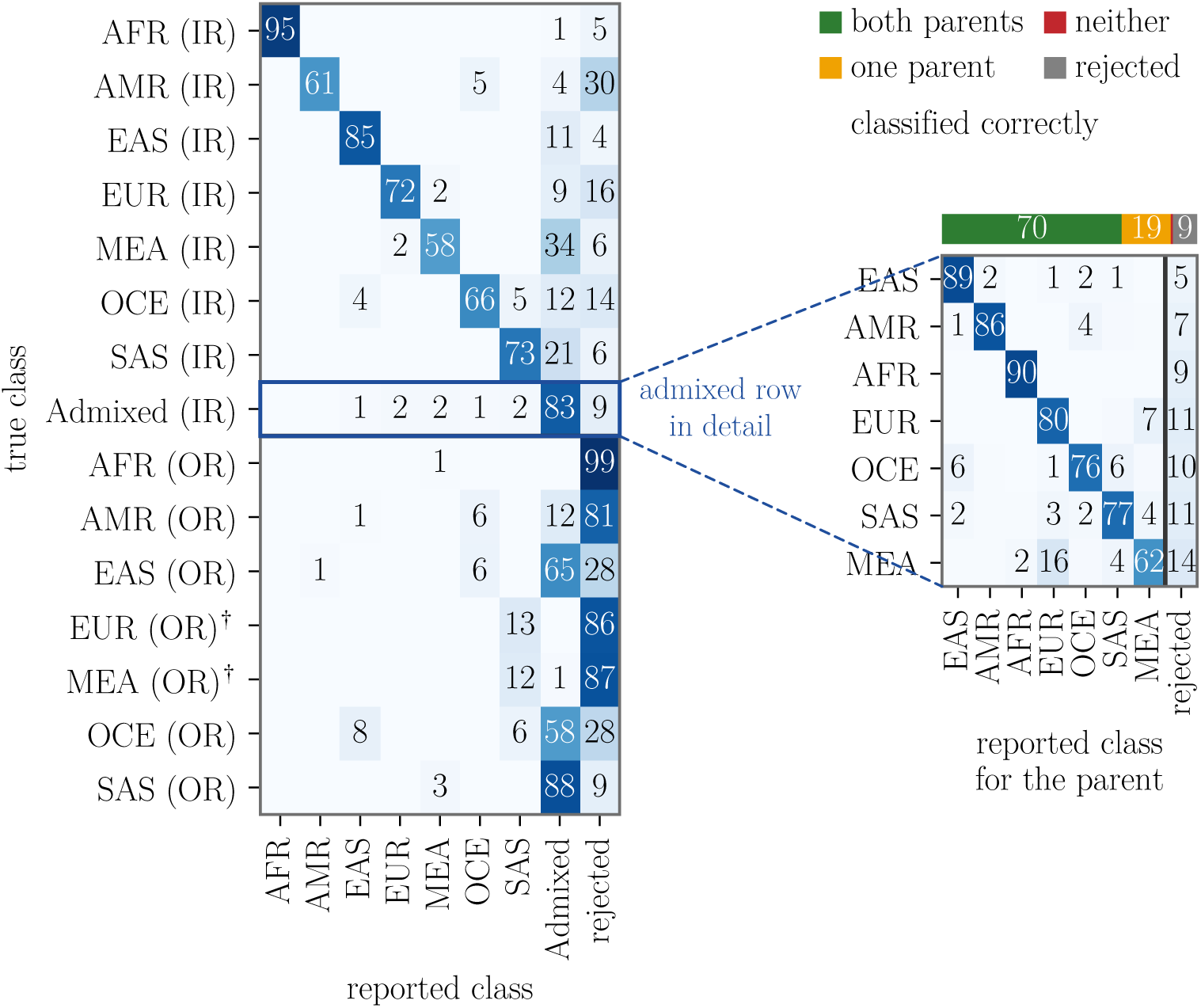
The admixture model under *s*_lik_. Among out-of-reference individuals, 0.370 are assigned to the Admixed class rather than rejected. This reflects the flexibility of the admixture model, which can represent a genotype as a combination of reference populations even when none provides a sufficient fit on its own. The model recovers both parental populations in 0.704 of cases, the highest proportion among the four methods that provide parental information, and rejects 0.095, the lowest proportion among them. Parental populations are defined as the two populations with the largest fitted proportions rather than as an explicitly modelled pair, which explains the similarity of the resulting matrix to those of the pair-based methods.

**Figure 11:**
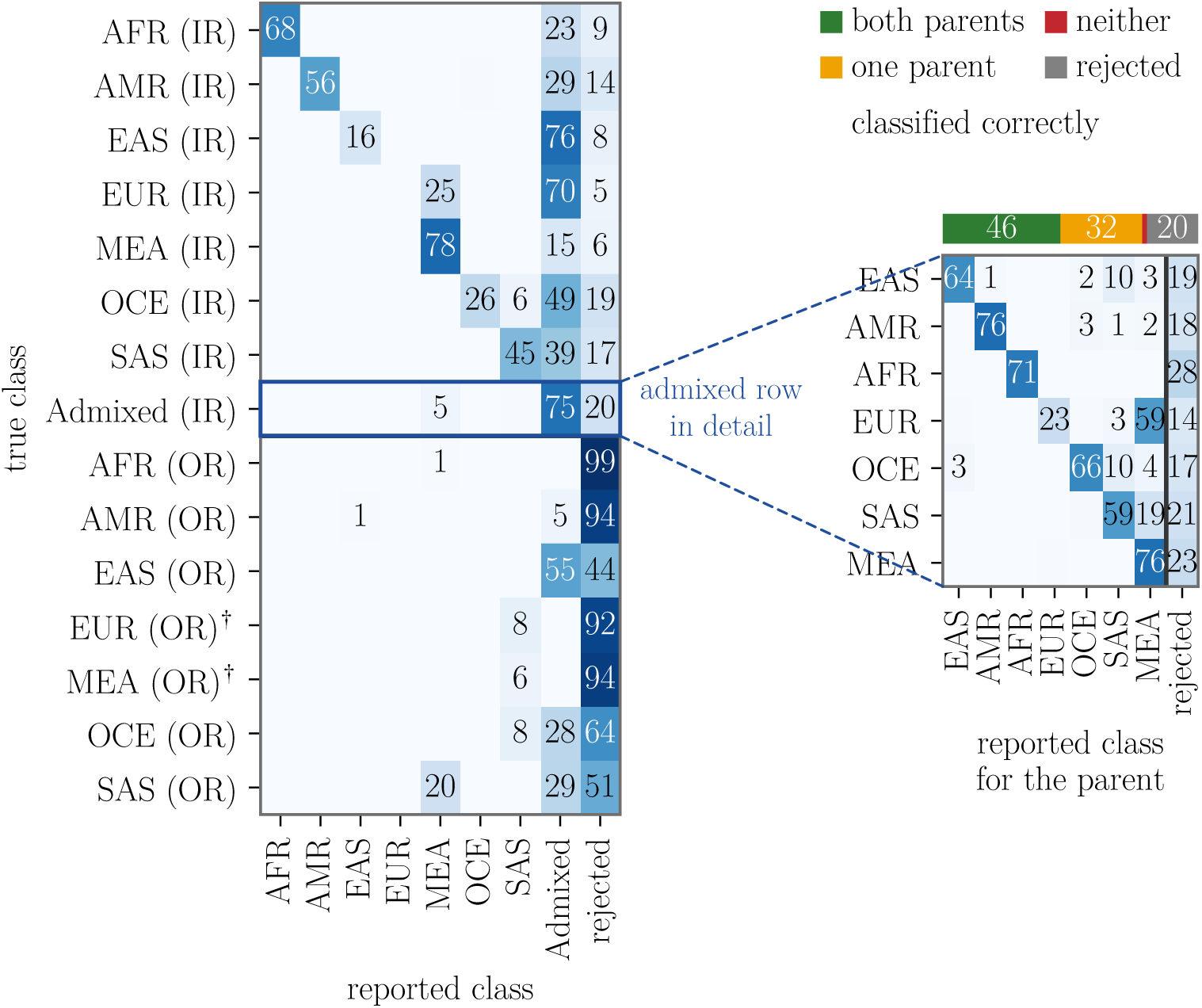
The pairwise goodness-of-fit statistic under either score. The method frequently assigns in-reference individuals to the Admixed class: 0.513 of truly in-reference individuals receive this label. Its in-reference rate of 0.332 therefore mainly reflects the class assignment made before application of the acceptance rule rather than the placement of the threshold. It is also the weakest of the four methods that provide parental information: both parental populations are recovered in 0.461 of cases, with 0.200 rejected. The loss is concentrated in EUR, the same population that drives the over-assignment to Admixed (fig. 5).

**Figure 12:**
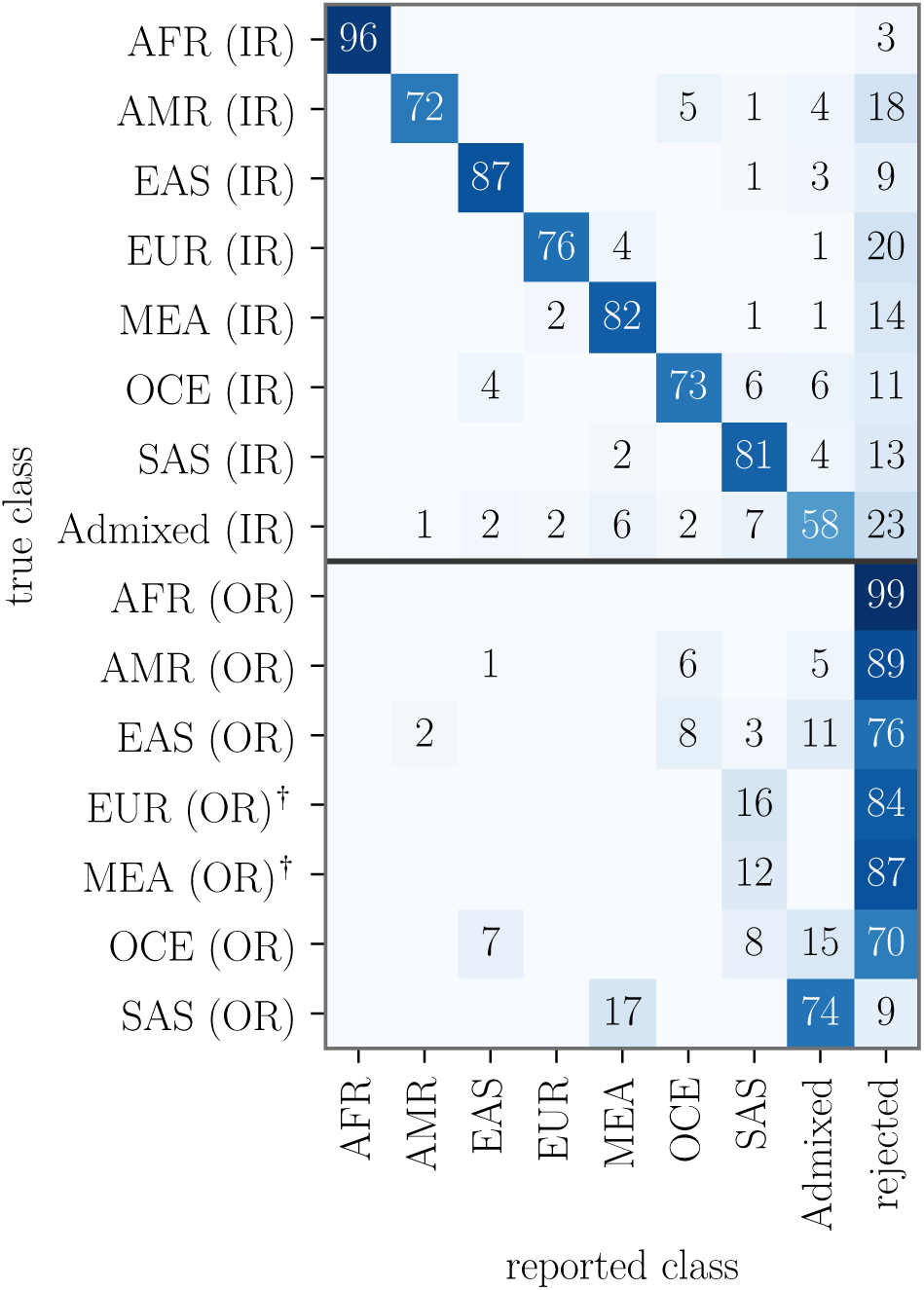
Naive Bayes over populations under *s*_lik_, after applying the class-specific acceptance thresholds. Errors occur in both directions: 0.192 of admixed individuals receive a population label, whereas 0.187 of out-of-reference individuals are labelled Admixed. This pattern results from the pooled Admixed class learnt from simulated offspring.

**Figure 13:**
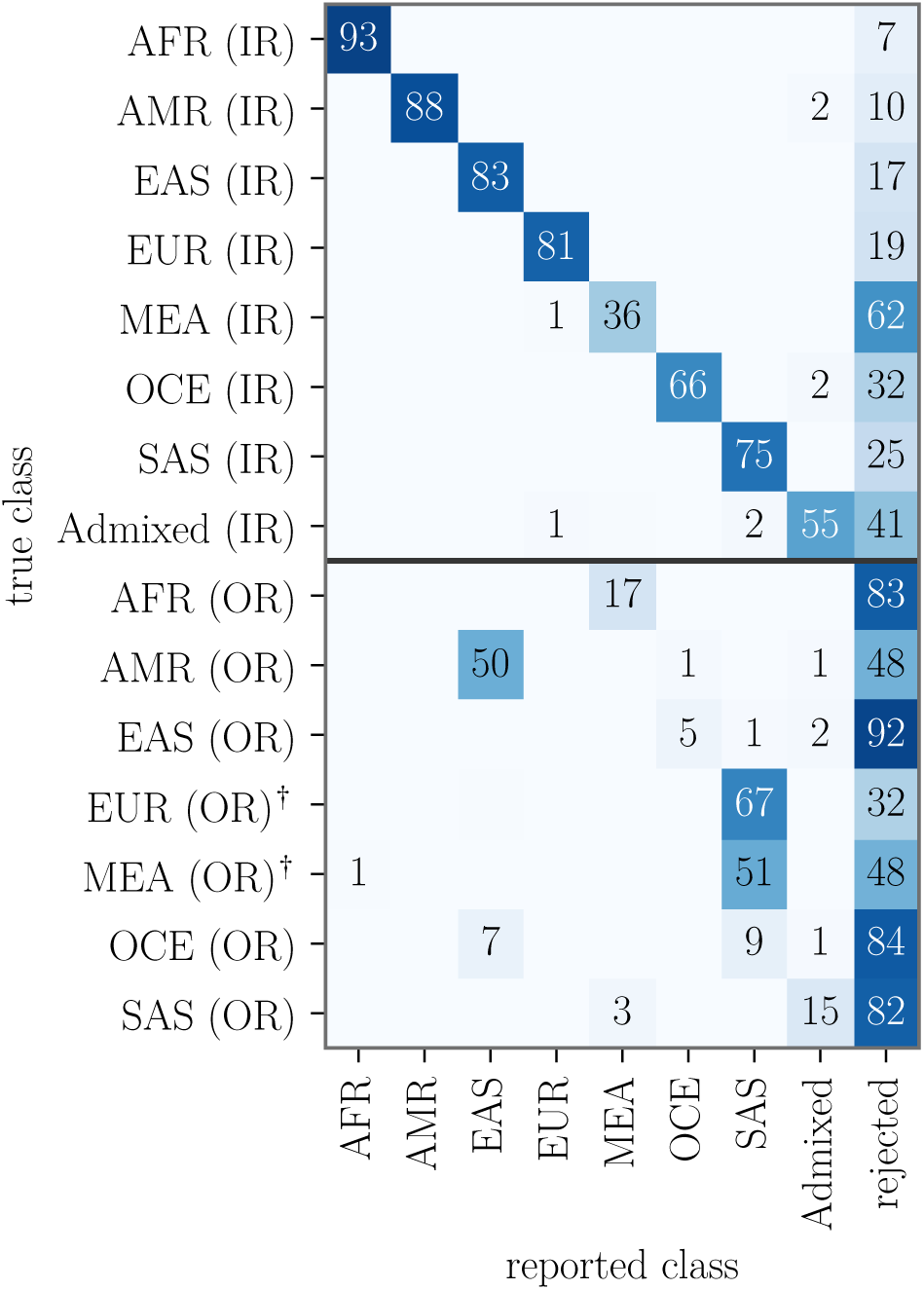
The tabular transformer under *s*_post_, after applying the class-specific acceptance thresholds. The largest loss occurs for admixed individuals, of whom 0.412 are rejected, the highest proportion among the methods shown here. Its posterior is normalised over the classes available during training, which can provide good relative ranking among those classes but cannot directly represent the possibility that none of them provides an adequate fit.

**Figure 14:**
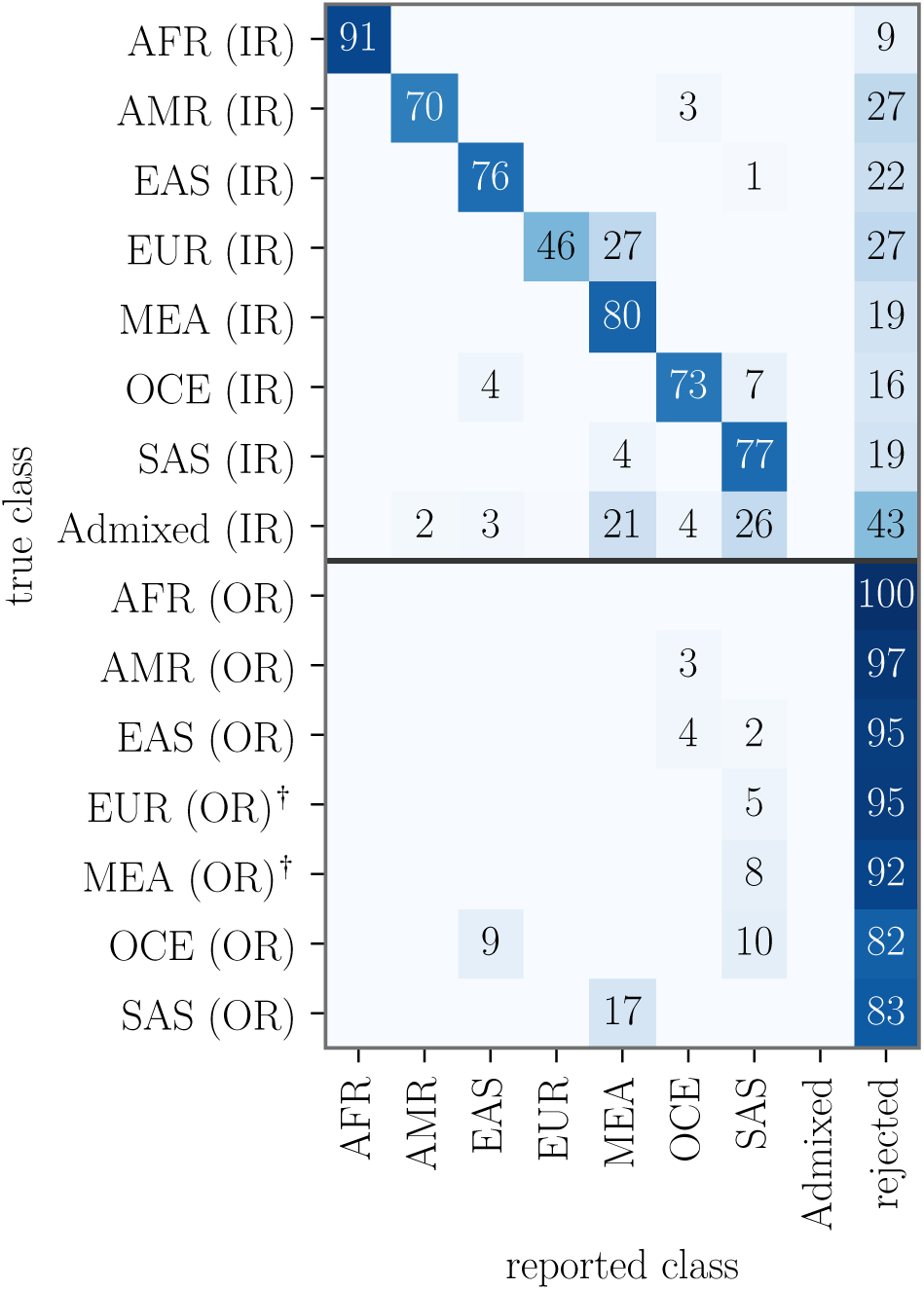
The single-population goodness-of-fit statistic, after applying the class-specific acceptance thresholds. The Admixed column is empty because the statistic does not represent two parental populations. At the same time, this method has the highest rejection rate among the six comparison methods, which explains why its AUC is highest in every design while its criterion value is zero. The figure therefore illustrates why both quantities are reported side by side.

### A.10 The 28 newly analysed samples, one row each

Table 5 lists the 28 samples of section 3.4 individually. The *stated ancestry* records the ancestry reported by the donor for each parent, including the reported country or region. The *target* maps this information to the classes represented in the reference panel and therefore provides the operational reference against which the prediction is compared. *Markers* gives the number of the panel’s 104 markers available for the respective sample. One marker, rs12594144, failed in almost every sample, and two additional markers of the assay are not part of the panel, so the attainable maximum is 103. The acceptance score *s*_lik_ is compared with the threshold of the predicted class. The margin is defined as their difference, so a sample is retained when the margin is non-negative.

### A.11 The remaining groups of the reference table, group by group

Table 2 in section 3.3 summarizes all seven groups; here, we describe the groups individually. Among the 47 east african individuals, which form the group with the largest distance from the reference panel and have a rejection rate of 0.83, 37 are described as a mixture of AFR and MEA in the underlying pair assignment. The roma group contains only three individuals, so it has little influence on the aggregate results and its *F_ST_*estimate should be interpreted with caution. The 136 (sanger) middle east individuals represent the same broad region as the panel’s MEA population but originate from a different data source (Almarri et al., 2021). Their distance to the panel’s MEA population is *F_ST_* = 0.007, and 0.25 are rejected. This group therefore provides information on the behaviour of the MEA threshold for independently sourced individuals from a represented region rather than on rejection of an absent population. For a north african individual, MEA is the closest available reference population. Whether such an assignment should be counted as an error is therefore primarily a question of the reference panel’s definition and resolution rather than of the classification method.

The two admixed cohorts are included for completeness, although a unique correct classification is not defined for them. The classifier assigns the Admixed class to 0.61 of these individuals before application of the acceptance rule, while the acceptance rule rejects 0.76. This pattern suggests that the cohorts contain ancestry components not represented by the reference panel.

## B Likelihoods and tests

### B.1 Likelihoods of the compared methods

Assume we have unphased genetic training/reference data (*x_ikjm_*)*_i,k,j,m_*, where *x_ikjm_* ∈ {0, 1, 2} is the number of alleles *j* in individual *i* from population *k* at marker *m*. Because humans are diploid, 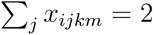 for all *i, k, m*, and 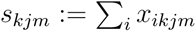 is the number of appearances of allele *j* in population *k* at marker *m*, such that 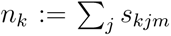 does not depend on *m* and is the number of (haploid) training samples from population *k*.

In addition, let (*y_m_*)*_m_* denote the test data, where *y_m_* = (*y_m_*_1_*, y_m_*_2_) is the pair of alleles observed at marker *m*. We describe the methods used in this study through their log-likelihoods, up to an additive constant, which are then maximized. Naive Bayes and Recent Naive Bayes yield classifications into one population or a pair of populations, respectively. Admixture and Recent Admixture instead estimate admixture proportions, denoted by *q*, for the test sample.

#### Naive Bayes

Assuming independence of the alleles in *y*,

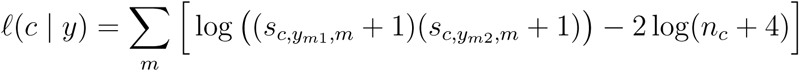

#### Recent Naive Bayes

For *k* ≠ *l*, either *y_m_*_1_ originates from population *k* and *y_m_*_2_ from population *l*, or vice versa, giving

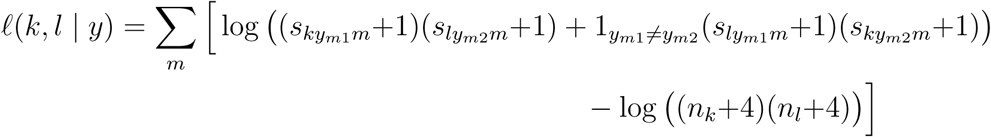

#### Admixture

Here, we set 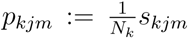 as the frequency of allele *j* in population *k* at marker *m*. Let *q* = (*q*_1_*, . . . , q_K_*), with *q_k_* ≥ 0 and 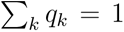, denote the proportions of the test individual’s genome originating from populations 1*, . . . , K*, i.e. the admixture proportions. Then,

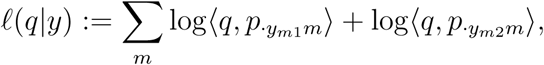

where 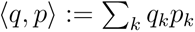 is the scalar product and *p_·jm_* := (*p_kjm_*)*_k_*_=1*,…,K*_.

#### Recent admixture

Let *q^′^* and *q^′′^* denote the maternal and paternal admixture proportions of the individual, respectively. The corresponding log-likelihood is

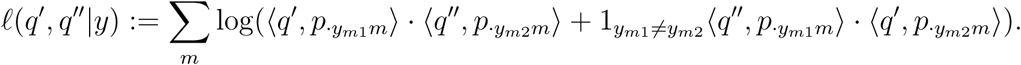

### B.2 Tests from Tvedebrink

Tvedebrink et al. (2018) considered the following test problem for a given population *k*:

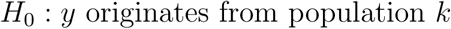

against the alternative

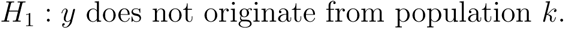

For the biallelic markers considered by Tvedebrink et al. (2018), only the counts *s_k_*_0*m*_ and *s_k_*_1*m*_ are required.

We define *s*_+*km*_ := *y_m_*+ *s_k_*_1*m*_ as the total count of allele 1 at locus *m* in population *k* together with the test individual. A likelihood-ratio approach for biallelic markers leads to the following locus-specific statistic. Writing 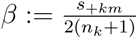 for the allele frequency in population and individual together and 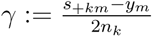 for the one in the population alone,

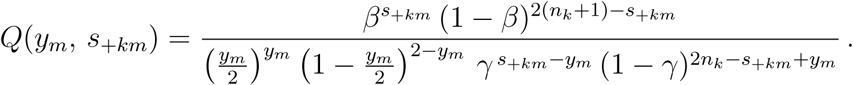

Assuming independence across markers, the resulting test statistic is

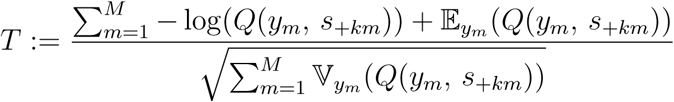

which is assumed to follow a normal distribution. To calculate the expectation and variance in this expression, we use

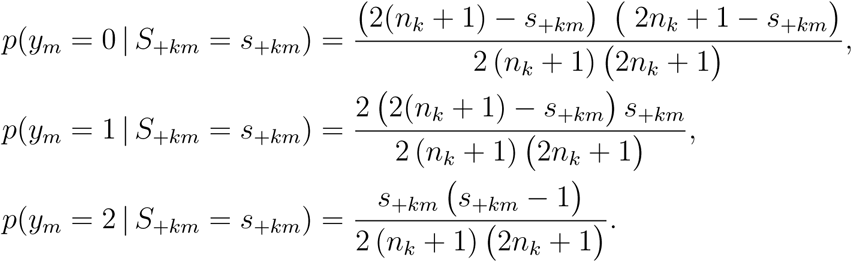

#### Multiallelic markers

Tvedebrink et al. (2018) restrict the analysis to biallelic markers, whereas the panel used here also contains multiallelic markers. We therefore treat each allele *a* of a marker as a separate locus, “*a* against the remaining 2 − *a* copies”, and sum the numerator and variance in *T* over these allele-specific loci in the same way as the original statistic sums over markers. One allele per marker is redundant and is dropped as the reference, so a bi-allelic marker leaves a single allele and *T* reduces to the original statistic. Only fully typed loci enter, and the reference panel is built on the training fold, so an allele unseen there is treated as missing rather than paired against a count of zero.

#### TVP: the pairwise extension

Tvedebrink and Eriksen (2019) extend the test to a pair of populations. Under

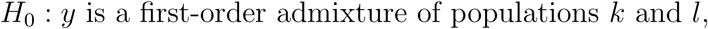

one allele of the individual comes from each parent, so with *p*^*_kam_* = (*s_kam_* + 1)*/*(2*n_km_* + 2) the Laplace-smoothed frequency of allele *a* at marker *m* in population *k*, and *y_am_* ∈ {0, 1, 2} its count in the profile,

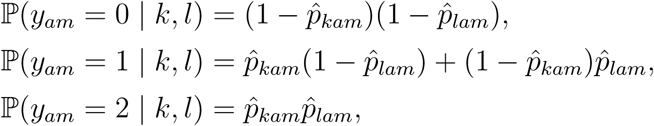

which is the same convolution used by the RNB classifier. Against the alternative of a random draw at allele frequency 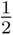 , that is 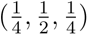 as in Tvedebrink and Eriksen (2019), the log-likelihood ratio per locus is log *Q_am_* = log P(*y_am_* | *k, l*) − log P(*y_am_* | *H*_1_), and the pair is scored by

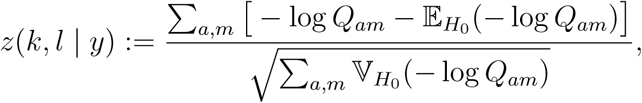

the sums running over the typed loci of the individual. Mean and variance are taken exactly over *y_am_* ∈ {0, 1, 2} under *H*_0_ rather than by the delta approximation of Tvedebrink and Eriksen (2019); *z* is then approximately standard normal when (*k, l*) is the true pair, and larger values of *z* provide stronger evidence against the pair. The pair (*k, k*) is the single-population test above, so the 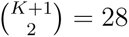 candidate pairs contain the *K* = 7 single populations as their diagonal. The individual is assigned the pair attaining the smallest *z*. This is reported as a single population when *k* = *l* and as Admixed otherwise. The quantity min*_k,l_ z*(*k, l* | *y*) is used for the acceptance rule, with its sign inverted because smaller values indicate a better fit.

Compared with Tvedebrink and Eriksen (2019), we make one simplification: the profile is not added back to the reference counts, i.e. we omit the EM step for allocating heterozygous alleles to the two parents. This affects the calibration of *z* but not the ranking of the population pairs.

## Notes

### Competing Interest Statement

The authors have declared no competing interest.

## References

Alexander, D. H., J. Novembre, and K. Lange (2009). Fast model-based estimation of ancestry in unrelated individuals. Genome research 19 (9), 1655–1664.

Almarri, M. A., M. Haber, R. A. Lootah, P. Hallast, S. Al Turki, H. C. Martin, Y. Xue, and C. Tyler-Smith (2021). The genomic history of the middle east. Cell 184 (18), 4612–4625.

Balding, D. J. and R. A. Nichols (1995). A method for quantifying differentiation between populations at multi-allelic loci and its implications for investigating identity and paternity. Genetica 96, 3–12.

Bartlett, P. L. and M. H. Wegkamp (2008). Classification with a reject option using a hinge loss. Journal of Machine Learning Research 9, 1823–1840.

Berk, R., L. Brown, A. Buja, K. Zhang, and L. Zhao (2013). Valid post-selection inference. The Annals of Statistics 41 (2), 802–837.

Bhatia, G., N. Patterson, S. Sankararaman, and A. L. Price (2013). Estimating and interpreting *F_ST_* : the impact of rare variants. Genome Research 23 (9), 1514–1521.

Butler, K., M. Peck, J. Hart, M. Schanfield, and D. Podini (2011). Molecular “eyewitness”: forensic prediction of phenotype and ancestry. Forensic Science International: Genetics Supplement Series 3 (1), e498.

Cawley, G. C. and N. L. C. Talbot (2010). On over-fitting in model selection and subsequent selection bias in performance evaluation. Journal of Machine Learning Research 11, 2079–2107.

Chow, C. K. (1970). On optimum recognition error and reject tradeoff. IEEE Transactions on Information Theory 16 (1), 41–46.

Cortes, C., G. DeSalvo, and M. Mohri (2016). Learning with rejection. In Algorithmic Learning Theory (ALT 2016), Volume 9925 of Lecture Notes in Computer Science, pp. 67–82. Springer.

de la Puente, M., L. Casanova-Adán, J. González-Bao, J. Pardo-Seco, A. Mosquera-Miguel, A. Ambroa-Conde, J. Ruiz-Ramírez, A. Cabrejas-Olalla, M. Boullón-Cassau, A. Freire- Aradas, A. Rodríguez, C. Phillips, and M. V. Lareu (2025). Evaluating the effect of marker panel sizes on estimation of bio-geographical co-ancestry proportions. Forensic Science International: Genetics 78, 103275.

de la Puente, M., J. Ruiz-Ramírez, A. Ambroa-Conde, C. Xavier, J. Pardo-Seco, J. Álvarez- Dios, A. Freire-Aradas, A. Mosquera-Miguel, T. E. Gross, E. Y. Y. Cheung, W. Branicki, M. Nothnagel, W. Parson, P. M. Schneider, M. Kayser, Á. Carracedo, M. V. Lareu, C. Phillips, et al. (2021). Development and evaluation of the ancestry informative marker panel of the visage basic tool. Genes 12 (8), 1284.

Dette, H., C. S. Heinzel, Z. Lange, and P. Pfaffelhuber (2026). Testing for single-population ancestry in the admixture model. arXiv preprint arXiv:2606.01990.

Diepenbroek, M., B. Bayer, K. Schwender, R. Schiller, J. Lim, R. Lagacé, and K. Anslinger (2020). Evaluation of the ion ampliseq phenotrivium panel: Mps-based assay for ancestry and phenotype predictions challenged by casework samples. Genes 11 (12), 1398.

Eduardoff, M., T. E. Gross, C. Santos, M. de la Puente, D. Ballard, C. Strobl, C. Børsting, N. Morling, L. Fusco, C. Hussing, B. Egyed, L. Souto, J. Uacyisrael, D. Syndercombe Court, Á. Carracedo, M. V. Lareu, P. M. Schneider, W. Parson, C. Phillips, et al. (2016). Interlaboratory evaluation of the euroforgen global ancestry-informative snp panel by massively parallel sequencing using the ion pgm. Forensic Science International: Genetics 23, 178–189.

Escher, L. M., M. S. Naslavsky, M. O. Scliar, Y. A. O. Duarte, M. Zatz, K. Nunes, and S. F. Oliveira (2022). Challenges in selecting admixture models and marker sets to infer genetic ancestry in a brazilian admixed population. Scientific Reports 12 (1), 21240.

Esposito, C., G. A. Landrum, N. Schneider, N. Stiefl, and S. Riniker (2021). Ghost: adjusting the decision threshold to handle imbalanced data in machine learning. Journal of Chemical Information and Modeling 61 (6), 2623–2640.

Fleckhaus, J., K. Anslinger, S. Lutz-Bonengel, M. Vennemann, and M. Diepenbroek (2025). Aktueller Stand der “erweiterten DNA-Analyse”. Rechtsmedizin 35, 405–413.

Freire-Aradas, A., C. Phillips, and M. V. Lareu (2017). Forensic individual age estimation with dna: From initial approaches to methylation tests. Forensic Science Review 29 (2), 121–144.

Garnier-Géré, P. and L. Chikhi (2013). Population subdivision, Hardy–Weinberg equilibrium and the Wahlund effect. In Encyclopedia of Life Sciences (eLS). Wiley. Covers both directions: the heterozygote deficit of a pooled structured sample, and the heterozygote excess of first-generation offspring of two differentiated populations.

Geifman, Y. and R. El-Yaniv (2017). Selective classification for deep neural networks. Advances in neural information processing systems 30.

Geng, C., S.-J. Huang, and S. Chen (2021). Recent advances in open set recognition: A survey. IEEE Transactions on Pattern Analysis and Machine Intelligence 43 (10), 3614–3631.

Heinzel, C. S., L. Purucker, F. Hutter, and P. Pfaffelhuber (2025). Advancing biogeographical ancestry predictions through machine learning. Forensic Science International: Genetics, 103290.

Hendrickx, K., L. Perini, D. Van der Plas, W. Meert, and J. Davis (2024). Machine learning with a reject option: A survey. Machine Learning 113 (5), 3073–3110.

Hendrycks, D. and K. Gimpel (2017). A baseline for detecting misclassified and out-of- distribution examples in neural networks. In International Conference on Learning Representations.

Herbei, R. and M. H. Wegkamp (2006). Classification with reject option. The Canadian Journal of Statistics 34 (4), 709–721.

Hollmann, N., S. Müller, K. Eggensperger, and F. Hutter (2023). TabPFN: A transformer that solves small tabular classification problems in a second. In Proceedings of the 11th International Conference on Learning Representations (ICLR).

Hollmann, N., S. Müller, L. Purucker, A. Krishnakumar, M. Körfer, S. B. Hoo, R. T. Schirrmeister, and F. Hutter (2025). Accurate predictions on small data with a tabular foundation model. Nature 637 (8045), 319–326.

Hudson, R. R., M. Slatkin, and W. P. Maddison (1992). Estimation of levels of gene flow from DNA sequence data. Genetics 132 (2), 583–589.

Kayser, M. (2015). Forensic dna phenotyping: Predicting human appearance from crime scene material for investigative purposes. Forensic Science International: Genetics 18, 33–48.

Kayser, M., W. Branicki, W. Parson, and C. Phillips (2023). Recent advances in forensic dna phenotyping of appearance, ancestry and age. Forensic Science International: Genetics 65, 102870.

Kayser, M. and P. de Knijff (2011). Improving human forensics through advances in genetics, genomics and molecular biology. Nature Reviews Genetics 12 (3), 179–192.

Kidd, K. K., W. C. Speed, A. J. Pakstis, M. R. Furtado, R. Fang, A. Madbouly, M. Maiers, M. Middha, F. R. Friedlaender, and J. R. Kidd (2014). Progress toward an efficient panel of snps for ancestry inference. Forensic Science International: Genetics 10, 23–32.

Kosoy, R., R. Nassir, C. Tian, P. A. White, L. M. Butler, G. Silva, R. Kittles, M. E. Alarcon-Riquelme, P. K. Gregersen, J. W. Belmont, F. M. De La Vega, and M. F. Seldin (2009). Ancestry informative marker sets for determining continental origin and admixture proportions in common populations in america. Human Mutation 30 (1), 69–78.

Kriegeskorte, N., W. K. Simmons, P. S. F. Bellgowan, and C. I. Baker (2009). Circular analysis in systems neuroscience: the dangers of double dipping. Nature Neuroscience 12 (5), 535–540.

Masana, M., I. Ruiz, J. Serrat, J. van de Weijer, and A. M. Lopez (2018). Metric learning for novelty and anomaly detection. arXiv preprint arXiv:1808.05492 .

McNevin, D. (2020). Forensic inference of biogeographical ancestry from genotype: The genetic ancestry lab. Wiley Interdisciplinary Reviews: Forensic Science 2 (2), e1356.

Mogensen, H. S., T. Tvedebrink, C. Børsting, V. Pereira, and N. Morling (2020). Ancestry prediction efficiency of the software genogeographer using a z-score method and the ances-try informative markers in the precision id ancestry panel. Forensic Science International: Genetics 44, 102154.

Naue, J. (2023). Getting the chronological age out of dna: using insights of age-dependent dna methylation for forensic dna applications. Genes & Genomics 45 (10), 1239–1261.

Pereira, R., C. Phillips, N. Pinto, C. Santos, S. E. dos Santos, A. Amorim, Á. Carracedo, and L. Gusmão (2012). Straightforward inference of ancestry and admixture proportions through ancestry-informative insertion deletion multiplexing. PLoS One 7 (1), e29684.

Pereira, V., A. Freire-Aradas, D. Ballard, C. Børsting, V. Diez, P. Pruszkowska-Przybylska, J. Ribeiro, N. M. Achakzai, A. Aliferi, O. Bulbul, M. D. P. Carceles, S. Triki-Fendri, A. Rebai, D. S. Court, N. Morling, M. V. Lareu, Á. Carracedo, et al. (2019). Development and validation of the euroforgen name (north african and middle eastern) ancestry panel. Forensic Science International: Genetics 42, 260–267.

Pereira, V., R. Santangelo, C. Børsting, T. Tvedebrink, A. P. F. Almeida, E. F. Carvalho, N. Morling, and L. Gusmão (2020). Evaluation of the precision of ancestry inferences in south american admixed populations. Frontiers in Genetics 11, 966.

Pfaffelhuber, P., F. Grundner-Culemann, V. Lipphardt, and F. Baumdicker (2020). How to choose sets of ancestry informative markers: A supervised feature selection approach. Forensic Science International: Genetics 46, 102259.

Pfaffelhuber, P., E. Sester-Huss, F. Baumdicker, J. Naue, S. Lutz-Bonengel, and F. Staubach (2022). Inference of recent admixture using genotype data. Forensic Science International: Genetics 56, 102593.

Phillips, C. (2015). Forensic genetic analysis of bio-geographical ancestry. Forensic Science International: Genetics 18, 49–65.

Phillips, C. and M. de la Puente (2021). The analysis of ancestry with small-scale forensic panels of genetic markers. Emerging Topics in Life Sciences 5 (3), 443–453.

Phillips, C., A. Salas, J. Sanchez, M. Fondevila, A. Gómez-Tato, J. Alvarez-Dios, M. Calaza, M. C. de Cal, D. Ballard, M. V. Lareu, et al. (2007). Inferring ancestral origin using a single multiplex assay of ancestry-informative marker snps. Forensic Science International: Genetics 1 (3-4), 273–280.

Pritchard, J. K., M. Stephens, and P. Donnelly (2000). Inference of population structure using multilocus genotype data. Genetics 155 (2), 945–959.

Rajeevan, H., U. Soundararajan, A. J. Pakstis, and K. K. Kidd (2012). Introducing the forensic research/reference on genetics knowledge base, frog-kb. Investigative Genetics 3, 18.

Rajeevan, H., U. Soundararajan, A. J. Pakstis, and K. K. Kidd (2020). Frogancestrycalc: a standalone batch likelihood computation tool for ancestry inference panels catalogued in frog-kb. Forensic Science International: Genetics 46, 102237.

Resutik, P., S. Aeschbacher, M. Krützen, A. Kratzer, C. Haas, C. Phillips, and N. Arora (2023). Comparative evaluation of the maplex, precision id ancestry panel, and visage basic tool for biogeographical ancestry inference. Forensic Science International: Genetics 64, 102850.

Resutik, P., M. Gysi, T. Lemmin, A. Kratzer, C. Haas, and N. Arora (2026). From unsupervised selection of aims to likelihood-based bga inference: Challenges revealed by openadmixture and genogeographer. Forensic Science International: Genetics 83, 103433.

Ruiz-Ramírez, J., M. de la Puente, C. Xavier, A. Ambroa-Conde, J. Álvarez-Dios, A. FreireAradas, A. Mosquera-Miguel, A. Ralf, C. Amory, M. A. Katsara, T. Khellaf, M. Nothnagel, E. Y. Y. Cheung, T. E. Gross, P. M. Schneider, J. Uacyisrael, S. Oliveira, M. D. N. Klautau-Guimarães, C. Carvalho-Gontijo, E. Póspiech, W. Branicki, W. Parson, M. Kayser, A. Carracedo, M. V. Lareu, and C. Phillips (2023). Development and evaluations of the ancestry informative markers of the VISAGE Enhanced Tool for Appearance and Ancestry. Forensic Science International: Genetics 64, 102853.

Samuel, G. and B. Prainsack (2019). Forensic dna phenotyping in europe: views “on the ground” from those who have a professional stake in the technology. New Genetics and Society 38 (2), 119–141.

Santos, C., C. Phillips, A. Gomez-Tato, J. Álvarez-Dios, Á. Carracedo, and M. V. Lareu (2016). Inference of ancestry in forensic analysis ii: Analysis of genetic data. In *Methods in Molecular Biology*, Volume 1420, pp. 255–285.

Scheirer, W. J., A. d. R. Rocha, A. Sapkota, and T. E. Boult (2013). Toward open set recognition. IEEE Transactions on Pattern Analysis and Machine Intelligence 35 (7), 1757–1772.

Schneider, P. M., B. Prainsack, and M. Kayser (2019). The use of forensic dna phenotyping in predicting appearance and biogeographic ancestry. Deutsches Árzteblatt International 116 (51-52), 873–880.

Shriver, M. D., M. W. Smith, L. Jin, A. Marcini, J. M. Akey, R. Deka, and R. E. Ferrell (1997). Ethnic-affiliation estimation by use of population-specific dna markers. American Journal of Human Genetics 60 (4), 957–964.

Smyser, T. J., P. Pfaffelhuber, R. M. Giglio, M. G. DeSaix, A. J. Davis, C. F. Bowden, M. A. Tabak, A. Manunza, V. A. Bâlteanu, M. Amills, et al. (2024). Probabilistic genetic identification of wild boar hybridization to support control of invasive wild pigs (sus scrofa). Ecosphere 15 (2), e4774.

Terrado-Ortunõ, N. and P. May (2024). Forensic dna phenotyping: a review on snp panels, genotyping techniques, and prediction models. Forensic Sciences Research 10 (1), owae013.

Tillmar, A. and D. Kling (2025). Comparative study of statistical approaches and snp panels to infer distant relationships in forensic genetics. Genes 16 (2), 114.

Truelsen, D., T. Tvedebrink, H. S. Mogensen, M. S. Farzad, M. A. Shan, N. Morling, V. Pereira, and C. Børsting (2021). Assessment of the effectiveness of the euroforgen name and precision id ancestry panel markers for ancestry investigations. Scientific Reports 11 (1), 18595.

Tvedebrink, T. and P. S. Eriksen (2019). Inference of admixed ancestry with ancestry informative markers. Forensic Science International: Genetics 42, 147–153.

Tvedebrink, T., P. S. Eriksen, H. S. Mogensen, and N. Morling (2017). Genogeographer – a tool for genogeographic inference. Forensic Science International: Genetics Supplement Series 6, e463–e465.

Tvedebrink, T., P. S. Eriksen, H. S. Mogensen, and N. Morling (2018). Weight of the evidence of genetic investigations of ancestry informative markers. Theoretical Population Biology 120, 1–10.

Varma, S. and R. Simon (2006). Bias in error estimation when using cross-validation for model selection. BMC Bioinformatics 7, 91.

Vidaki, A. and M. Kayser (2018). Recent progress, methods and perspectives in forensic epigenetics. Forensic Science International: Genetics 37, 180–195.

Vovk, V., A. Gammerman, and G. Shafer (2005). Algorithmic Learning in a Random World. New York: Springer.

Wang, S., C. M. Lewis, M. Jakobsson, S. Ramachandran, N. Ray, G. Bedoya, W. Rojas, M. V. Parra, J. A. Molina, C. Gallo, G. Mazzotti, G. Poletti, K. Hill, A. M. Hurtado, D. Labuda, W. Klitz, R. Barrantes, M. C. Bortolini, F. M. Salzano, M. L. Petzl-Erler, L. T. Tsuneto, E. Llop, F. Rothhammer, L. Excoffier, M. W. Feldman, N. A. Rosenberg, and A. Ruiz-Linares (2007). Genetic variation and population structure in native americans. PLoS Genetics 3 (11), e185.

Wu, M. and J. Ye (2009). A small sphere and large margin approach for novelty detection using training data with outliers. IEEE transactions on pattern analysis and machine intelligence 31 (11), 2088–2092.

Xavier, C., M. de la Puente, C. Phillips, M. Eduardoff, A. Heidegger, A. Mosquera-Miguel, A. Freire-Aradas, R. Lagace, S. Wootton, D. Power, W. Parson, M. V. Lareu, and R. Daniel (2020). Forensic evaluation of the asia pacific ancestry-informative maplex assay. Forensic Science International: Genetics 48, 102344.

Zieger, M. (2022). Forensic dna phenotyping in europe: How far may it go? Journal of Law and the Biosciences 9 (2), lsac024.

